# Metagenomic domain substitution for the high-throughput modification of non-ribosomal peptide analogues

**DOI:** 10.1101/2023.05.31.543161

**Authors:** Sarah R. Messenger, Edward M. R. McGuinniety, Luke J. Stevenson, Jeremy G. Owen, Gregory L. Challis, David F. Ackerley, Mark J. Calcott

## Abstract

Non-ribosomal peptides are a diverse and medically important group of natural products. They are biosynthesised by modular non-ribosomal peptide synthetase (NRPS) assembly-lines in which domains from each module act in concert to incorporate a specific amino acid into a peptide. This modular biosynthesis has driven efforts to generate new peptide analogues by substituting amino acid specifying domains. Rational NRPS engineering has increasingly focused on using evolutionarily favoured recombination sites for domain substitution. Here, we present an alternative approach inspired by evolution, which involves large-scale diversification and screening. By adopting a metagenomic approach of amplifying amino acid specifying domains from metagenomic DNA derived from soil, we were able to substitute over 1,000 unique domains into a pyoverdine NRPS. To identify functional domain substitutions, we employed fluorescence and mass spectrometry screening techniques, followed by sequencing. This comprehensive screening process successfully identified more than 100 functional domain substitutions, resulting in the production of 16 distinct pyoverdines as major products. The significance of this metagenomic approach lies in its ability to shift the focus of engineering non-ribosomal peptide biosynthesis. Instead of relying on a high success rate of individual domain substitution, we have developed effective methods that enable the exploration of a broader range of substitutions. This opens new possibilities for the discovery and production of novel non-ribosomal peptides with diverse biological activities.

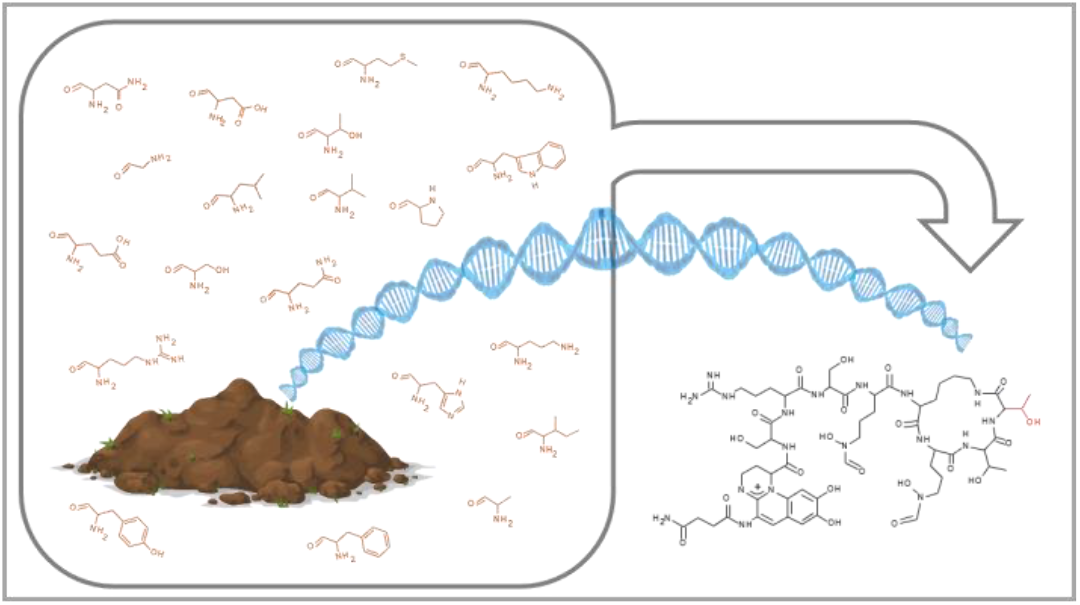

## Introduction

More than half of approved therapeutic drugs are natural products or derived from natural products.^1^ Discovery of their therapeutic potential has been a linchpin of modern medicine. For example, antibiotics such as penicillin have allowed effective treatment of infections and reduced the risk of life-threatening infections for other medical interventions like surgeries.^2^ Penicillin falls within the non-ribosomal peptide class of natural product, an essential group of compounds for human health that includes antibiotics, anticancer drugs and immunosuppressants.^3^ Non-ribosomal peptides can have complex structures that include features that are necessary for bioactivity or bioavailability, but are very difficult to synthesise chemically,^4,5^ e.g. large sizes with many stereocentres, methylation of amide bonds, non-proteinogenic amino acids and strained ring systems. The relative cost and efficiency of fermentation means major pharmaceutical companies often rely on bacterial fermentation to produce non-ribosomal peptide drugs, for example bleomycin, ciclosporin, colistin and vancomycin; all of which are considered essential medicines by the World Health Organisation.^6^

While natural evolution has provided a plethora of chemical diversity, non-ribosomal peptides have not evolved to function optimally in the human body. The requirement for production by fermentation limits the capacity to improve current drugs or modify compounds with undesirable properties to be suitable therapeutics. Due to increasing rediscovery rates as more compounds are known, extensive efforts to find and isolate new or better drugs from nature have met diminishing returns.^7^ These factors have driven synthetic biology-based approaches to create microbes that produce better drugs. Examples of such approaches are finding gene clusters to heterologously express from metagenomic DNA,^8-11^ or engineering the enzymes within known biosynthetic pathways to produce improved analogues.^5,12-14^

NRPS enzymes are particularly appealing targets for engineering due to their modular assembly-line architecture. Each NRPS module within an assembly-line recognises and incorporates a specific amino acid onto a growing peptide (Fig. 1a). This mode of biosynthesis offers scope for creating new peptides by substituting domains of individual enzyme modules with others that cause a different amino acid to be added at a particular location (Fig. 1b). This has proven difficult to achieve in the lab, with early work that emphasised keeping paired C-A domains intact generally having low success rates and product yields.^12-14^ More recently, progress has been made exploring new recombination sites^15-20^ with an expanding emphasis being to understand and target evolutionarily favoured recombination sites.^15,18,21-23^ We previously found this was useful for identifying structurally favoured sites for entire A domain substitutions.^15^ At the same time, our evolutionary analyses indicated that partial A domain substitutions occur more frequently in nature. However, partial A domain substitutions appear more prone to unfavourable structural disruptions, and we found these to be more difficult than entire A domain substitutions to recapitulate effectively in the laboratory setting. An explanation for the prevalence of partial A domain substitutions when examining evolutionary events is that these substitutions occur more frequently in nature.^15^ Collectively, these observations spurred us to investigate whether inefficiencies in experimental domain substitution can be overcome by developing a method for large scale diversification and selection.

**Fig. 1:**
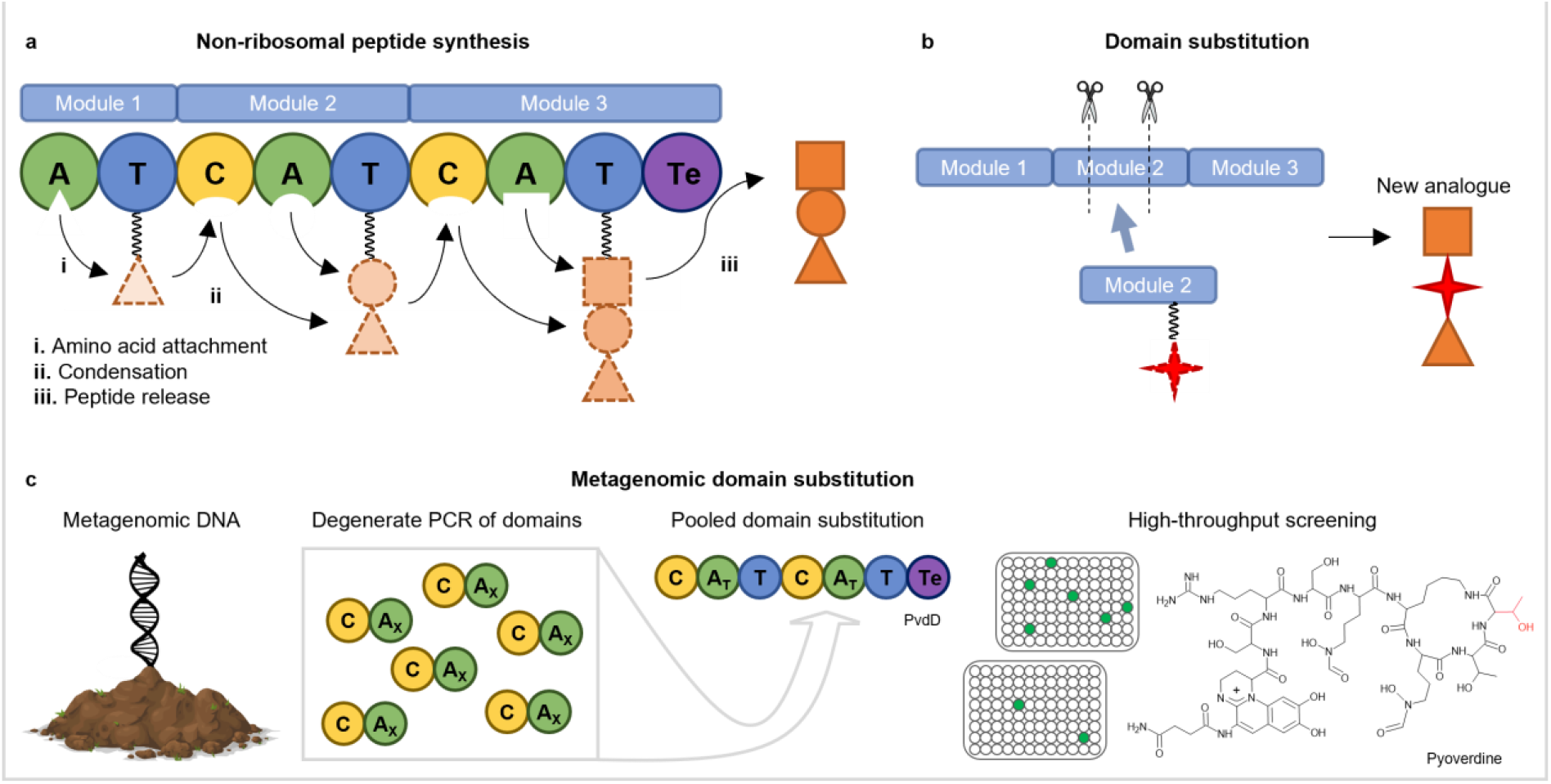
Non-ribosomal peptide synthesis and metagenomic domain substitution. **a**. During non-ribosomal peptide synthesis, an amino acid substrate is adenylated by an adenylation (A) domain and attached to a phosphopantetheine cofactor linked to the downstream thiolation (T) domain. Condensation (C) domains then catalyse amide bond formation between substrates attached to adjacent modules. The growing peptide is passed down the NRPS assembly line until it is released by a thioesterase (Te) domain or alternative release domain within the final module. **B**. Domain substitution aims to create analogues by replacing amino acid specifying domain(s) with domains that incorporate alternative amino acids. **C**. Metagenomic domain substitution uses pools of unknown domains amplified from environmental DNA for substitution. Screening is then performed to identify bacterial strains producing modified non-ribosomal peptides. This study used metagenomic domain substitution to modify the final amino acid of pyoverdine (indicated in red) by replacing domains in the second module of PvdD, a two module NRPS that (in its native form) incorporates two L-Thr residues at the *N*-terminal end of pyoverdine. AT refers to Thr-specific A domains; AX refers to A domains that have specificity for unknown amino acids. The terminal L-Thr residue modified in this study is highlighted in red.

To source domains for large scale diversification, we turned our attention to soil metagenomic DNA. Efforts to discover new biosynthetic pathways have shown that metagenomic DNA derived from soil is rich in NRPS genes. In particular, next-generation sequencing of A domains amplified using degenerate primers showed that a single extraction of total soil DNA can contain many thousands of unique A domains.^24,25^ This suggested that it might be possible to perform substitutions at scale by amplifying conserved domain regions from unknown NRPS within metagenomic DNA and using the resulting pools of domains for *en masse* domain substitution experiments.

Here we show that conserved motifs within NRPS genes can be suitable recombination sites for domain substitution, and identify preferred locations to substitute diverse pools of domains into module 2 of *pvdD*, the final module of pyoverdine biosynthesis in *Pseudomonas aeruginosa* PAO1 (Fig. 1c). This allowed us to perform the parallel insertion of over a thousand unique domains into *pvdD*. By transforming the resulting libraries into *P. aeruginosa* PAO1 and screening for pyoverdine producing bacteria, we successfully exemplified large scale domain substitution and screening as a successful strategy for generating non-ribosomal peptide diversity.

## Results

### Selection of highly conserved sequences to use as recombination boundaries

We first aimed to identify candidate recombination sites within highly conserved regions of DNA that would maximise amplification from metagenomic DNA using degenerate primers. There are known to be seven conserved amino acid motifs within C domains (C1-C7) and ten within A domains (A1-A10).^26^ As most of these exhibit substantial variation at a DNA level, it was unclear which regions would be optimal primer-binding sites. We analysed DNA sequence diversity using previously-compiled^15^ alignments of 650 NRPS modules from two biosynthetically rich genera^27^ with similar GC content (437 modules from *Pseudomonas* and 213 from *Streptomyces*). Information present, a metric of diversity based on Shannon entropy,^28^ was calculated at each position of the alignments. This was averaged over 18 base segments (Fig. 2) and indicated that the most conserved DNA sequences were indeed centred on the conserved motifs, although the A1 and A4 motifs were relatively weakly conserved. Additional conserved regions were also identified between the C2 & C3, and C7 & A1 motifs. The two conserved regions between the C7 & A1 motifs (labelled 1 and 2 in Fig. 2) were upstream to the C-terminus of the C-A domain linker (labelled X in Fig. 2), a region we previously found successful for A domain substitution.^15^ Overall, this sequence analysis highlighted multiple regions of highly conserved DNA that might serve as sites for designing degenerate primer pairs. Moreover, the consistent pattern of sequence conservation between the genera supported that primers could be designed that would amplify domains from distantly related genome templates.

**Fig. 2:**
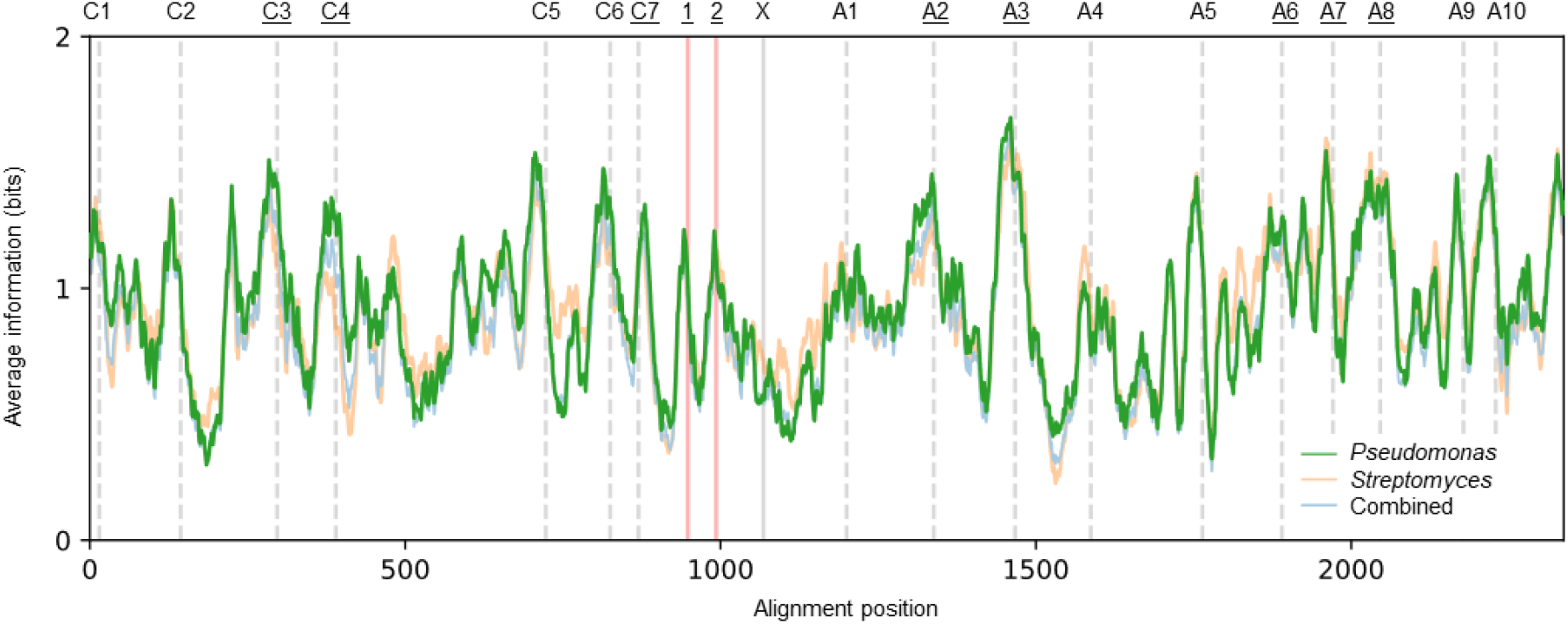
Identification of candidate recombination sites based on sequence conservation. Diversity (information present)^28^ averaged over 18 bp fragments along the lengths of sequence alignments comprising 437 *Pseudomonas* and 213 *Streptomyces* C-A domains. The locations of the C1-C7 and A1-A10 motifs are indicated by vertical grey dashed lines. Two conserved locations between the C7 and A1 motifs are highlighted by vertical red solid lines, and a site upstream to the C-A domain linker is labelled ‘X’ and shown by a vertical solid grey line. The labels at the top of the graph for motifs and other recombination sites that were selected for testing in subsequent experiments are underlined.

We next sought to identify promising primer-binding sites that would also minimise structural perturbations when used as recombination sites for domain substitution. We considered this would likely increase the proportion of functional recombination events. The structural assessment was first performed using SCHEMA, a tool to predict amino acid perturbations introduced at each possible recombination site between two proteins.^29^ Using the crystal structure of the C-A domains from the first module of the teixobactin synthesising enzyme Txo2 (pdb: 6P1J),^30^ we created a profile of the number of perturbations predicted to be introduced for every recombination site. This compared sites between module two of PvdD, and each of eight modules we knew to be capable of functional substitutions into PvdD using the C1-A10 motifs as recombination boundaries. Two of these eight modules, specifying the amino acids Ser or *N*^5^-formyl-*N*^5^-hydroxyornithine (fhOrn) were previously used for functional substitutions into Pa11 as either C-A domain pairs or A-domains.^15^ The remaining six, specifying Gly, Ala, Arg, Arg, Glu or Phe, were found to be functional during preliminary testing of metagenomic domain library building using the C1-A10 motifs (Supplementary Fig. 1). The profile indicated that recombination at the following motif positions would cause the fewest perturbations: C3, C4, halfway between the C7 and A1, and the continuous sequence from A6 to A10 (Fig. 3a). We then used ThreaDomEx as an additional tool to identify boundaries likely to be tolerant of domain substitution. ThreaDomEx uses threading alignments of multiple PDB files to predict the boundaries of structurally conserved subunits that can fold and evolve independently from each other.^31^ This analysis yielded partially congruent results to the SCHEMA analysis, with boundaries for conserved subunits predicted to be at the C4, halfway between the C7 and A1, and at the A8 motifs (Fig. 3b). The predictions appeared sensible when rationalised against the crystal structure 6P1J used for the SCHEMA analysis. In particular, they were all located between domains or subdomains and therefore more likely to keep the core structure intact (Fig. 3c).

**Fig. 3:**
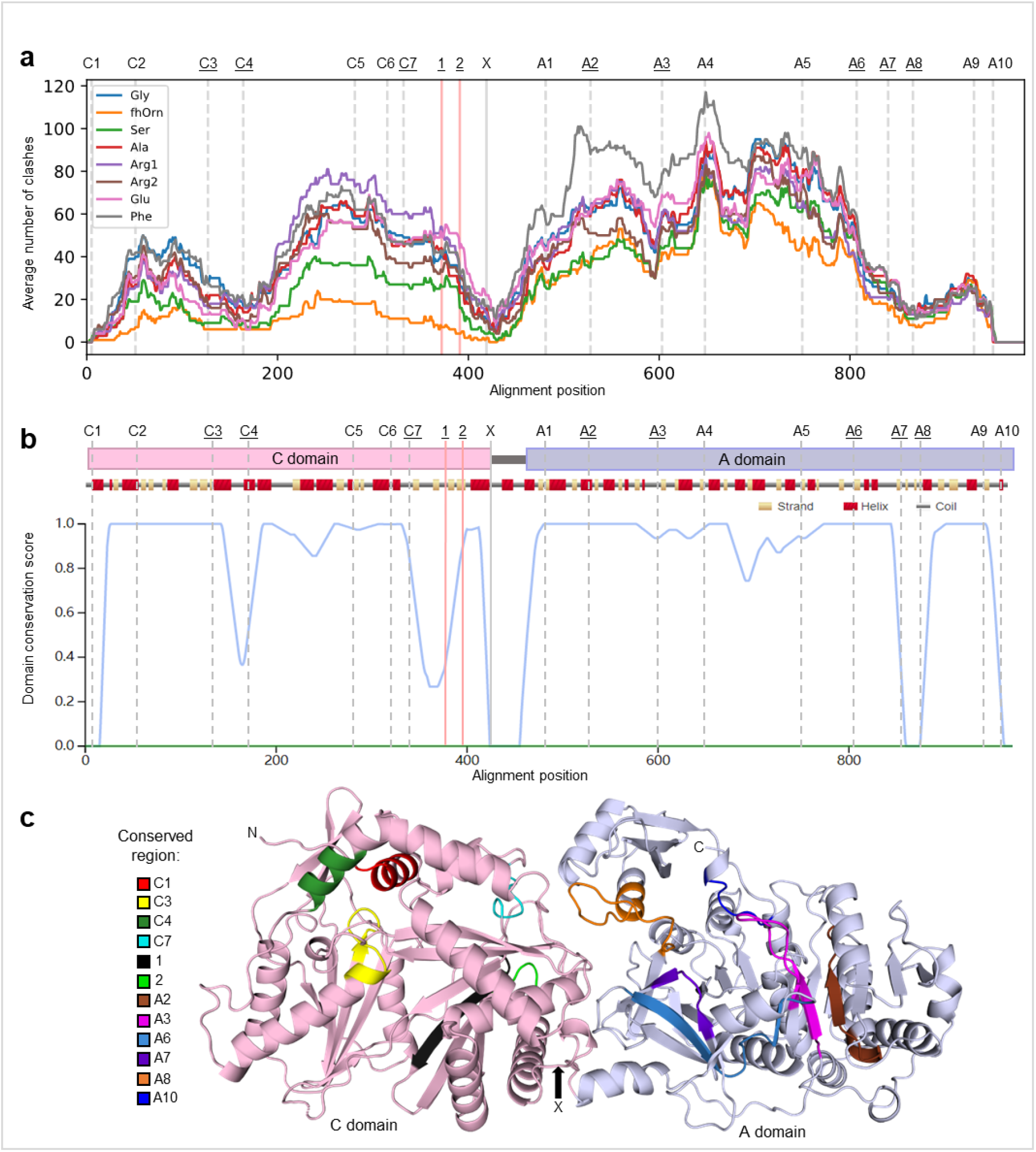
Structural analysis of candidate recombination sites. **a**. SCHEMA profile showing the predicted number of clashes introduced at each amino acid position when modelling recombination of the second module of PvdD with each of eight alternative modules (identities and rationale for selection explained in Supplementary Fig. 1). Within the inset key, standard abbreviation codes are used to identify the amino acids encoded by each alternative module, with fhOrn designating *N*^5^-formyl-*N*^5^-hydroxyornithine. The labelling of motif and candidate recombination sites is as per Fig. 2. **b**. The domain conservation profile produced for the second module of PvdD using ThreaDomEx, which assigns low conservation scores to sequences predicted to border regions of independent folding. Labelling is the same as panel a. **c**. The crystal structure of 6P1J with highlighted regions identified as candidate recombination sites that were tested in this work.

Based on the sequence and structural analyses, we selected candidate recombination sites to experimentally test in our pyoverdine model. Alternative domains were cloned into the corresponding region of the second module of a plasmid-based copy of *pvdD* and then used to transform a *P. aeruginosa pvdD* deletion strain followed by testing of pyoverdine production. Potential upstream and downstream recombination sites were first tested independently. The upstream sites were located at the C3, C4, C7, A2 and A3 motifs as well as two sites between the C7 and A1 motifs. While the A2 and A3 sites appeared structurally unfavourable (Fig. 3, panels a and b), they were selected due to being hotspots for partial A domain recombination in nature.^15^ A first set of substitutions were constructed using the same Gly, fhOrn and Ser modules employed for the SCHEMA profile in Fig. 3a. These substitutions were made by varying the upstream recombination site while keeping the downstream site at the A10 motif. As these substitutions were preselected to be functional substitutions using a downstream A10 site, this allowed us to examine the relative efficiency of each candidate upstream site. We observed that the C4-A10 domain substitutions were particularly well tolerated, with all three *P. aeruginosa* PAO1 strains containing these recombinant PvdD enzymes producing the expected pyoverdine analogue (Supplementary Fig. 2), and two yielding a statistically significant increase in pyoverdine production relative to substitutions at the C1 motif (Fig. 4a). Candidate downstream recombination sites were then tested in a similar fashion, with the upstream recombination site being upstream of the C-A domain linker (labelled X in Figs. 2, 3 and 4),^15^ while the downstream recombination site was varied. Candidate sites were tested within the A6, A7 and A8 motifs, with a total of three alternative sites tested within the A8 motif due to its longer length as well as both SCHEMA and ThreaDomEx suggesting it would be particularly tolerant of substitution. This time, each new candidate recombination site performed worse than the A10 motif in terms of pyoverdine yield (Fig. 4b).

**Fig. 4:**
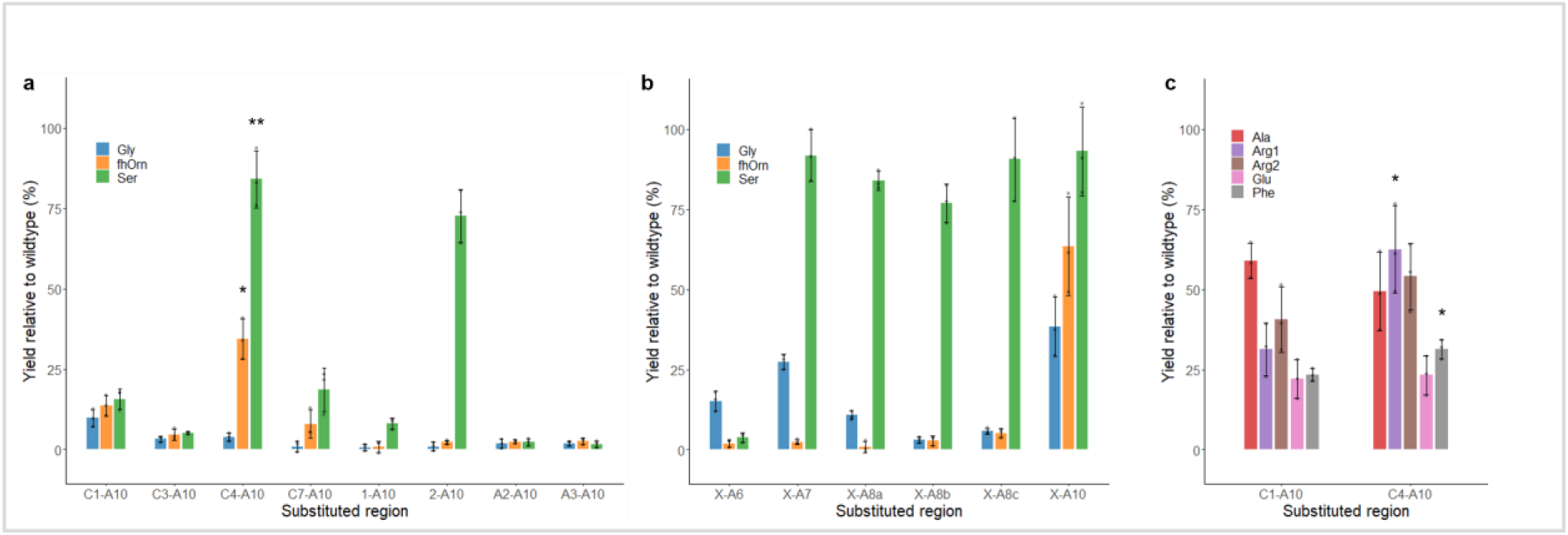
Experimental testing of candidate recombination sites. **a**. Pyoverdine yield for substitution variants that varied the upstream recombination site and maintained a constant tolerant downstream recombination point within the A10 motif. **B**. Pyoverdine yield for substitution variants that varied the downstream recombination site and maintained a constant tolerant upstream recombination site immediately upstream to the C-A domain linker. **c**. Pyoverdine yield comparing additional C1-A10 and C4-A10 domain substitutions. In all cases, pyoverdine yield was assessed by measuring absorbance at 400 nm relative to a wild-type *P. aeruginosa* strain grown under identical conditions. Mass spectrometry data is provided for successful C1-A10 and C4-A10 domain substitutions in Supplementary Fig. 2. All recombination sites are labelled as per Figs. 2 and 3, with the corresponding sequences indicated in Supplementary Fig. 1. Sites A8a to A8c refer to three distinct recombination sites tested within the A8 motif. Standard amino acid abbreviations are used, with Arg1 and Arg2 distinguishing two different Arg-specific modules. The abbreviation fhOrn represents *N*5-formyl-*N*5-hydroxyornithine. In all cases, n = 3 independent experiments and data are presented as mean values ± SD. Source data are provided as a Source Data file. Significantly improved for C4-A10 domain substitutions relative to the equivalent C1-A10 domain substitutions were calculated using an unpaired Welch two sample t-test. \**P* < 0.1; \*\**P* < 0.01.

As the C4-A10 domain substitutions appeared to be the best candidate recombination sites based on these initial substitutions (X-A10 substitutions not being an option as the X site had insufficient sequence conservation to permit degenerate primer design; Fig. 2), we next performed C4-A10 domain substitutions into the second module of PvdD using the other five modules identified in preliminary testing. Including the initial three substitutions, the C4-A10 substitution strains resulted in an increase in pyoverdine production in 4/8 cases relative to C1-A10 substitution strains, and there were similar levels of production for the remaining cases (Fig. 4, panels a and c). Given that the corresponding C1-A10 domain substitutions had been preselected for functionality, the increased pyoverdine production from C4-A10 domain substitution strains supported that the C4 and A10 motifs were superior candidate recombination sites to use for degenerate domain substitution.

### Degenerate primer design and PCR testing

Informed by the preliminary testing above, we sought to design degenerate primers to perform C4-A10 domain substitutions. Supporting this choice of recombination site, the C4 motif is located between two subdomains of the C domain and this recombination site,^17,32^ and a similar recombination site,^33^ have previously been tested in multiple pathways, and the A10 motif has likewise been tested in multiple pathways.^15,34,35^ As a point of comparison, we also designed primers that would enable C1-A10 domain substitutions because substitutions that keep partner C-A domain pairings intact have been used in many NRPS pathways.^12-14^ To identify primer sets with the greatest degeneracy and lowest non-specific amplification, we first used the tool HYDEN^36^ to design 24 sets of degenerate primers that differed in length, degeneracy, and binding region. These were designed to maximise amplification from the pool of *Pseudomonas* and *Streptomyces* sequences assessed in Fig. 2. *In silico* analysis showed that up to 40% of the *Pseudomonas* and *Streptomyces* DNA sequences contained primer binding sites, allowing for one mismatch per primer (Supplementary Table 1). Having been designed from a relatively high GC dataset, we considered these primer sets would likely bias toward GC-rich templates, and indeed, a far smaller proportion of the 370 *Bacillus* (lower GC) sequences we collated previously^15^ contained plausible binding sites (Supplementary Table 1).

Primer sets were experimentally tested for their relative abilities to amplify DNA from two metagenomic DNA libraries that we created from DNA extracted from soil and cloned into cosmids (Supplementary Table 2). The two libraries, referred to here as RX and SL, contained 8.5 and 12.9 million colony forming units respectively. Each library was arrayed across four 96-well plates with approximately 11,000 to 46,000 colony forming units per well (Supplementary Table 2). Preliminary testing involved amplification of samples from eight wells under the same PCR conditions for all 24 primer sets, with sets 13, 22 and 23 producing amplicons of the expected size for the C1-A10 region from the most samples (Supplementary Fig. 3). As primer sets 22 and 23 had higher degeneracy and were predicted to amplify more sequences than primer set 13 (Supplementary Table 1), we reasoned that they would provide the greatest breadth of amplification, and chose primer set 22 due to having the highest level of amplification (Supplementary Fig. 3). We next used HYDEN to design 12 forward primers around the C4 motif that possessed similar levels of degeneracy to primer set 22 (Supplementary Table 1). These forward primers were then tested in conjunction with the preferred degenerate reverse primer from primer set 22 and DNA template from three randomly chosen wells of the metagenomic libraries, followed by testing of the top performing primer with a further eight random samples (Supplementary Fig. 4). Based on this testing, forward primer 10 coupled with the reverse primer of primer set 22 was selected to amplify the C4-A10 domains for metagenomic domain substitution library construction.

### Construction of metagenomic domain substitution libraries

The C1-A10 domain substitution libraries were first created while we tested the alternative recombination sites described above. To do this, a plasmid was constructed with silent SpeI and NotI restriction sites bordering the C1 and A10 motifs within module 2 of *pvdD*. Equivalent restriction sites were added to the 5’ end of each degenerate primer and these were then used to amplify the C1-A10 regions from all eight 96-well plates of metagenomic DNA (Supplementary Fig. 5). The PCR products from each 96-well plate were pooled and DNA corresponding to the expected size of the C1-A10 regions was gel purified. The resulting pools of domains were then inserted into *pvdD* by restriction cloning. In generating these C1-A10 libraries, we noticed the ligation efficiency was low, which we considered might be due in part to inefficiencies introduced by the added restriction sites and associated spacer region in the primers. We therefore switched to Gibson assembly for creating the C4-A10 libraries, as this enabled us to use the originally-optimised primer sets. Following amplification and gel purification, Gibson assembly proceeded efficiently, resulting in the C4-A10 libraries being approximately 15-fold larger than the C1-A10 libraries in terms of colony forming units (Supplementary Table 3). In total, from the eight RX1-4 and SL1-4 metagenomic template plates, we created 16 sub-libraries comprised of eight for the C1-A10 domain substitutions and eight for the C4-A10 domain substitutions (Fig. 5a).

**Fig. 5:**
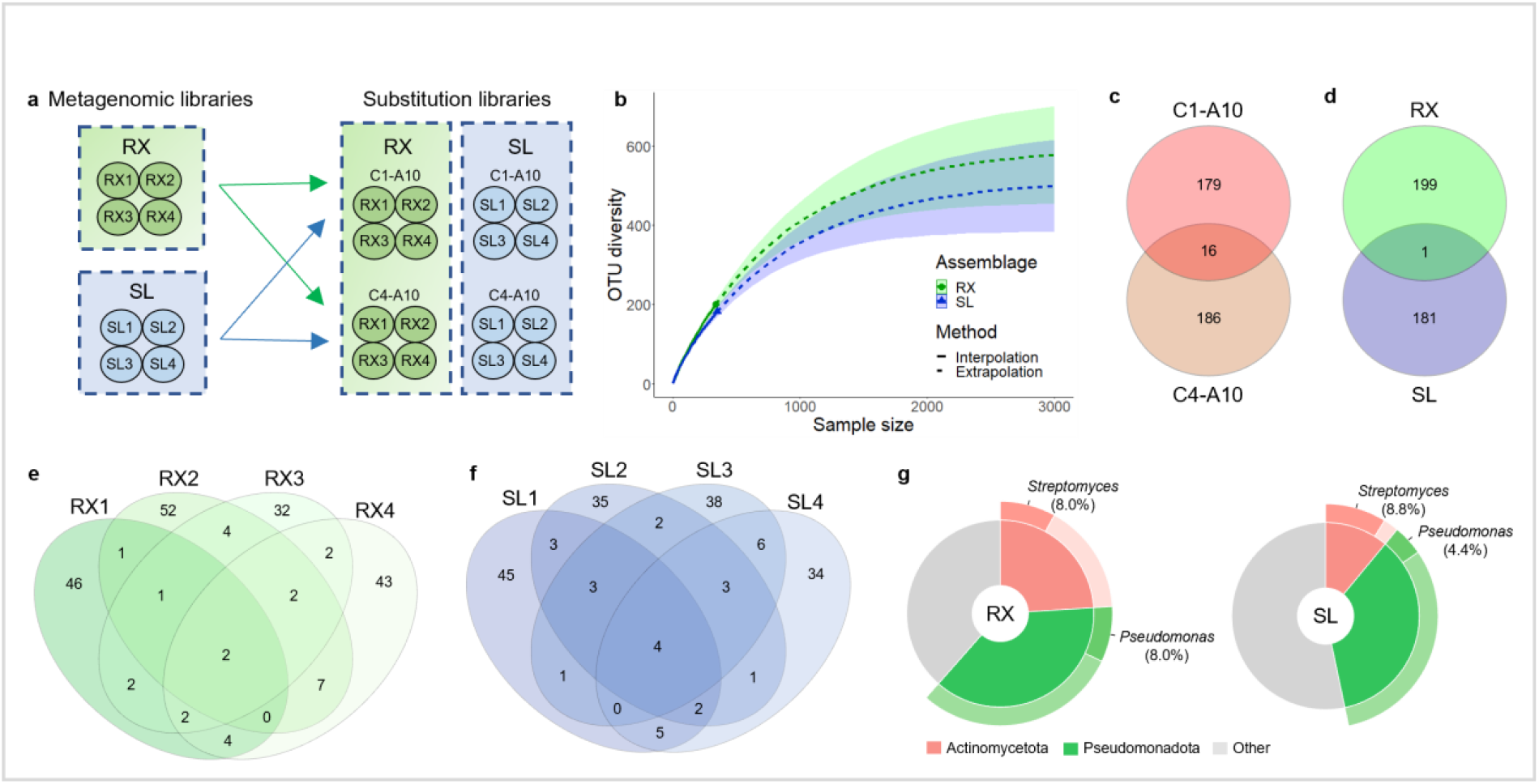
Diversity of the metagenomic domain substitution libraries. **a**. Schematic indicating how the RX and SL metagenomic cosmid libraries were used to derive the RX and SL substitution libraries. The RX and SL cosmid libraries were arrayed over four 96 well plates, and each plate was treated as a separate sub-library to create corresponding C1-A10 and C4-A10 substitution libraries. In total, this yielded an RX library comprising RX1-RX4 sub-libraries for each of the C1-A10 and C4-A10 domain substitutions, and an SL library comprising SL1-SL4 sub-libraries for each of the C1-A10 and C4-A10 domain substitutions. **b**. The OTU diversity of the RX and SL libraries calculated using the python package iNEXT.^37^ Solid lines represent estimates of OTU diversity based on interpolation of sequencing data. Dashed lines represent estimates of OTU diversity extrapolated from sequencing data. **c**. Venn diagram showing the number of distinct and shared OTUs in the C1-A10 and C4-A10 domain substitutions identified from 685 total sequences. **d**. Venn diagram showing the number of distinct and shared OTUs in the SL and RX libraries identified from 685 total sequences. **e**. Venn diagram showing the numbers of distinct and shared OTUs found in the RX1-RX4 sub-libraries. **f**. Venn diagram showing the numbers of distinct and shared OTUs found in the SL1-SL4 sub-libraries. **g**. The inner donut chart represents the proportion of OTUs for which the closest BLASTx match was from the phyla Actinomycetota or Pseudomonadota. The outer donut chart shows the percentage of OTUs that had a closest match to *Streptomyces* or *Pseudomonas*.

To examine the diversity of the C1-A10 and C4-A10 libraries, we sequenced a subset of samples and analysed the sequences. A total of 840 colonies including at least 48 colonies from each sub-library were randomly selected for plasmid purification, and the ends of each insert within module 2 of *pvdD* were Sanger sequenced in the forward and reverse direction. The reverse sequences were used to analyse diversity between the RX and SL libraries (chosen because these gave sequence spanning the same region for both C1-A10 and C4-A10 domain substitutions). Sequences were processed to ensure high quality reads, which resulted in 685 total sequences with 35-57 from each sub-library. Clustering the sequences at 95% identity into operational taxonomic units (OTUs) identified 381 unique OTUs. The R package iNEXT^37^ was then used to estimate the total number of OTUs in each library. This found an increase in new OTUs without reaching a plateau as the sequencing data set increased, and the total population size was estimated to be 597 OTUs for the RX library and 516 OTUs for the SL library (Fig. 5b). There was a low level of overlap in sequences found for C1-A10 and C4-A10 domain substitutions (Fig. 5c), and only one OTU was found in both the RX and SL library (Fig. 5d). A greater degree of overlap was identified between sub-libraries (Fig. 5, panels e and f), which was consistent with the sub-libraries being derived from the same original soil DNA. To provide an indication of bacterial diversity, each OTU was analysed by BLASTx and the highest sequence identity hit was assessed according to its taxonomic grouping. The top BLASTx hits were predicted to be from twelve phyla (Fig. 5g) with 54% from the same phyla as *Pseudomonas* and *Streptomyces*, i.e. Pseudomonadota and Actinomycetota. Greater diversity was present at the genus level, with the top blast hits being from 95 genera (see source data file). Overall, these results indicate that although our primer design strategy did bias for sequences related to *Pseudomonas* and *Streptomyces*, the metagenomic domain libraries were nevertheless large and diverse, comprising over 1,000 unique NRPS sequences and embodying a high level of bacterial diversity.

### Library screening and analysis

To screen for functional substitutions, the libraries were transformed into a *P. aeruginosa* PAO1 strain lacking *pvdD*. In preliminary screening of the C1-A10 domain substitutions, transformed colonies were grown in 96-well plates at a dilution of ∼5 colonies per well. In this manner, an estimated 8,000 colonies were tested for pyoverdine production using MALDI-TOF mass spectrometry. Whilst this identified 94 strains that produced 11 different pyoverdines as major products (Supplementary Figs. 7 and 8), this screening had a low hit rate of ∼1% and isolating individual producing strains from their screening pools was time-consuming. We considered this pilot approach was likely missing many hits, which would make it difficult to compare between libraries. As such, we altered the screening to directly compare an equal number of individually selected clones for each of the 16 sub-libraries. A total of 4,608 transformed colonies (288 per sub-library) was picked and these were used to inoculate individual wells of 96-well plates. We have previously found that measuring fluorescence emission saturates quickly but allows detection of low levels of pyoverdine directly from whole-cell cultures, whereas absorbance has a linear relationship with pyoverdine concentration but requires only the supernatant to be tested.^38^ Therefore, a multi-stage process was used to first screen a single replicate from each colony for fluorescence and then more accurately measure pyoverdine yield from selected colonies by absorbance and by MALDI-TOF mass spectrometry to detect unique pyoverdines and predict the amino acid substitution (Fig. 6a). Overall, this second screening pipeline identified that 487 of the 4,608 colonies produced pyoverdines, a much higher hit rate (>10%) than the preliminary screen. Proportionately, there were 3.3 fold more hits from the C4-A10 domain substitutions than the C1-A10 domain substitutions (73 *vs*. 260 for the RX library and 41 *vs*. 113 for the SL library). Fourteen unique pyoverdines were identified as major products and the pyoverdine yield for hits was generally low with a few high-yield outliers (Fig. 6b; Supplementary Fig. 9). With the exception of valine, which was biased by a large number of hits from the RX library C4-A10 domain substitutions (Fig. 6b), the distribution of pyoverdine yield per substituted amino acid was similar for both the RX and SL libraries (Fig. 6c).

**Fig. 6:**
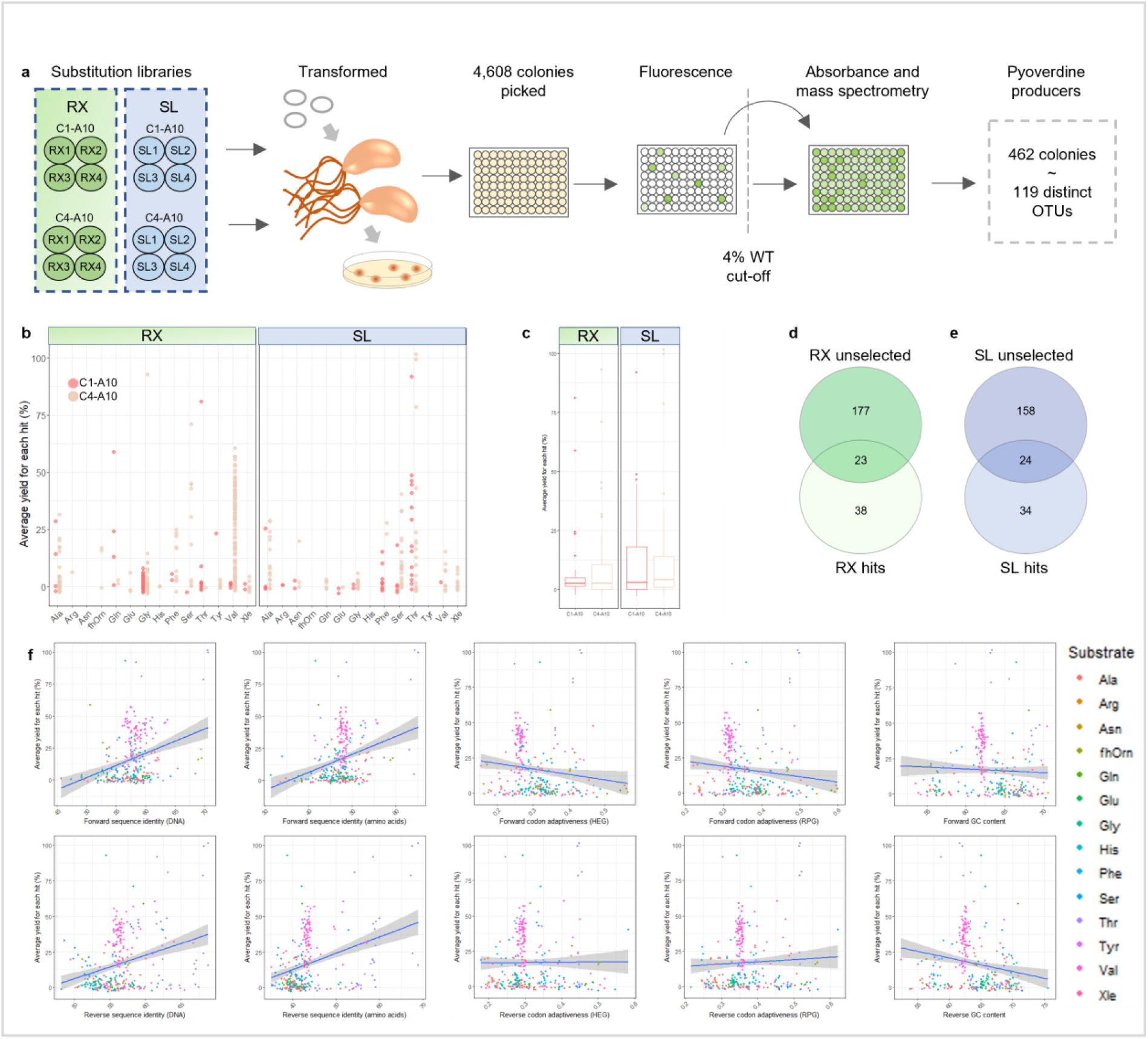
Screening and diversity of functional substitutions. **a**. Overview of the screening process to identify pyoverdine producing strains. **b**. The average pyoverdine yield for each hit. **c**. A boxplot showing average yield for all non-Val producing hits. **d & e**. Venn diagram showing the number of distinct OTUs from the unselected library colonies and the pyoverdine producing hits found during screening for the RX and SL libraries, respectively. **f**. Graphs showing average pyoverdine yield versus DNA sequence identity, amino acid identity, codon adaptiveness of highly expressed genes (HEG) in P. aeruginosa PAO1, codon adaptiveness of ribosomal protein genes (RPG) and GC content for the forward and reverse sequence of each insert. All data points represent at least three independent replicates.

To analyse the relationship between DNA sequence and function, the ends of the inserted domains were sequenced as above for the plasmids derived from 384 hits (4 × 96 well plates selected to exclude low yield Gly and Val substitutions due to the high number of hits for these substrates). Clustering the sequences at 95% identity gave 119 unique OTUs. Only 39% of these OTUs were also found in the OTUs from unselected colonies (Fig. 6, panel d and e). This confirms that the initial sequencing of unselected OTUs only represented a small portion of the total library size. Examining the relationship between pyoverdine production and either percent identity of the inserted domains to *pvdD*, codon-adaptiveness of the insert to *P. aeruginosa* PAO1, or GC content found weak or insignificant trends (Fig. 6f). The percent identity between the introduced and replaced domains had a weak positive relationship with product yield. However, this was skewed by a small number of Thr-specific domains that were closely related to the replaced region of *pvdD*. As these had the same specificity as the replaced domains, closely related substitutions such as these were not considered useful as they do not alter the amino acid added. Overall, this analysis suggests that sequence identity, codon usage and GC content were not major factors influencing the efficiency of metagenomic domain substitution.

The strain giving the highest yield for each substrate, for each of the C1-A10 and C4-A10 domain substitutions from both the RX and SL libraries, was selected for retesting in triplicate once to minimise the variation implicit in high-throughput measurements. When determining yield based on absorbance, the most effective C4-A10 domain substitutions tended to be higher yielding than the most effective C1-A10 domain substitutions (Fig. 7; Supplementary Fig. 9). As yield distributions did not vary markedly by amino acid substitution (Fig. 6c), this likely stemmed from the C4-A10 library yielding a greater proportion of functional hits. The highest yielding strains were further analysed by liquid chromatography mass spectrometry and their pyoverdines analysed using molecular networking (Supplementary Fig. 10, Supplementary Table 4). *P. aeruginosa* produces pyoverdine with a succinamide side-chain as the major product, which is known to be accompanied by minor products with succinic acid and α-ketoglutaric acid side-chain.^39,40^ Including two additional strains recovered only in the pilot screening, we identified sixteen strains producing different pyoverdine variants as major products. From these sixteen strains, LC-MS molecular networking was used to identify unique pyoverdines from each strain. Two major networks of pyoverdines were produced, one which contained the major products and side chain derivatives and a second network which were mostly pyoverdines chelating aluminium (a phenomenon previously noted by Cornelis^41^). His and Me-Glu pyoverdines were too low yield to be incorporated into the molecular network, however these were manually analysed to identify ions consistent with these substrates. In total, 40 unique pyoverdines were identified when including the minor products.

**Fig. 7:**
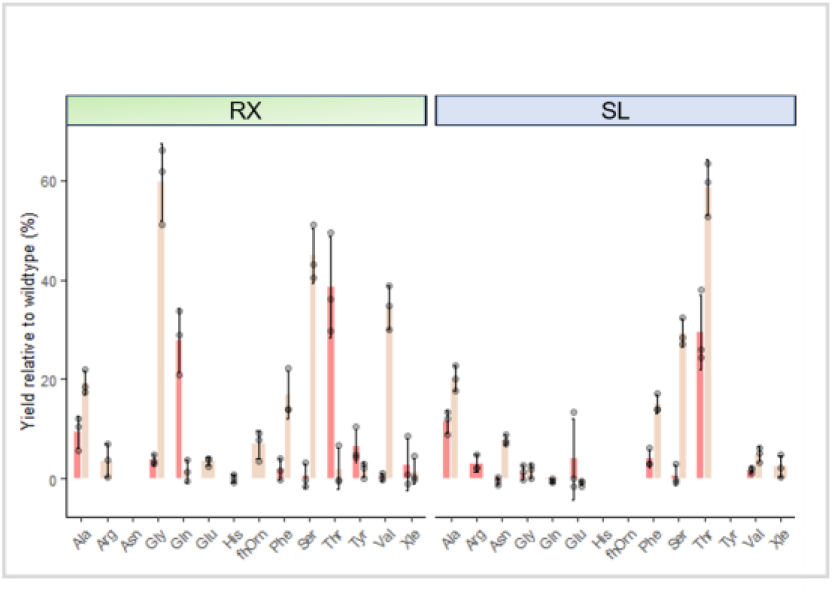
Pyoverdine yield fortop hits. **a**. Pyoverdine yield for the highest yielding hits from the RX and SL domain substitution libraries. The highest yielding hits were analysed for both C1-A10 and C4-A10 domain substitutions were analysed. In all cases, pyoverdine yield was assessed by measuring absorbance at 400 nm relative to a wild-type *P. aeruginosa* strain grown under identical conditions. Mass spectrometry data is provided for successful C1-A10 and C4-A10 domain substitutions in Supplementary Fig. 8 and 9. Standard amino acid abbreviations are used with fhOrn designating *N*^5^-formyl-*N*^5^-hydroxyornithine. In all cases, n = 3 independent experiments and data are presented as mean values ± SD. Source data are provided as a Source Data file.

## Discussion

We describe here a new high-throughput approach to domain substitution and its application to substitute over 1,000 domains into the second module of the pyoverdine synthetase PvdD. Screening the resulting domain substitution libraries allowed us to identify 119 successful functional domain substitutions, collectively yielding 16 unique pyoverdines as major products and a total of 40 pyoverdines overall. This number of substitutions into a single module is orders of magnitude greater than previous studies. For example, the largest previous report of domain substitution into a single module, that we are aware of, described the substitution of only twelve unique domains,^15^ and only combinatorial approaches of substitutions spanning multiple modules have approached comparable numbers of analogues produced.^14,18^ Although recent years have seen several new strategies implemented to improve the success rates of individual substitutions,^5,12-14,18^ a ‘one size fits all’ solution to NRPS engineering remains out of reach. We have instead sought a solution that is tolerant of the noisiness inherent in NRPS engineering. By developing a means to efficiently create large and diverse variant libraries, the requirement to maximise recombination efficacy diminishes and the problem instead becomes one of screening effectively.

Diversification followed by screening has been effective for engineering enzymes in other fields. Conceptually, degenerate domain substitution to target a specific residue in a non-ribosomal peptide is similar to saturation mutagenesis of codons for protein engineering, i.e. both approaches replacing a single encoded amino acid by numerous alternatives. Combined with screening, saturation mutagenesis has enabled probing of enzyme function and generation of focused directed evolution ‘smart libraries’ to engineer new or improved activities.^42,43^ Metagenomic domain substitution is the first example, to our knowledge, of performing such diversification with entire protein domains. Although many desirable non-ribosomal peptides lack a chromophore that would allow them to be screened spectrophotometrically like pyoverdine, other screening approaches have been utilised for decades by natural products researchers and may also be suitable for metagenomic domain substitution libraries. For instance, the MALDI-TOF mass spectrometry we describe here was particularly valuable and, along with LC-MS dereplication techniques,^44^ is likely applicable to other non-ribosomal peptides. High-throughput screens have already been used for discovering new NRPS pathways,^11^ restoring NRPS function,^45^ modifying A domain binding pockets,^46^ and screening for strains producing higher titres of non-ribosomal peptides.^47^ These studies show the potential for developing functional screens, which could allow domain substitution experiments to be performed in other NRPS pathways at a scale that is not possible with rational engineering. The missing piece of the puzzle – how to generate large domain substitution libraries – is something metagenomic domain substitution provides.

An additional important consideration for metagenomic domain substitution experiments is how to maximise the diversity generated. Our results show that degenerate primers can be used to efficiently amplify diverse NRPS domains from metagenomic DNA, targeting different conserved recombination points. The primers we designed were based on a comprehensive dataset of *Streptomyces* and *Pseudomonas* sequences, to enrich for domains from these biosynthetically gifted genera.^27^ It is likely that further diversity could be accessed by using complementary sets of primers that collectively provide broader specificity. Even so, our more-focused primers amplified over 1,000 unique domains and there was little overlap in DNA sequences identified from the RX and SL libraries we generated. This indicates that greater variation could have been extracted by increasing screening depth, and that more variation could undoubtedly be accessed with the same primers by targeting additional sources of metagenomic DNA.

A limitation of metagenomic domain substitution is that recombination boundaries are constrained to conserved motif sequences as recombination boundaries. This is different to rational NRPS strategies that have historically targeted loop regions that possess low sequence conservation. It is worth considering that natural NRPS diversification during natural evolution is most likely to occur via homologous recombination at conserved sequence regions, and we demonstrated here that multiple NRPS motif sequences are tolerant of recombination. Nevertheless, we note that our initial testing suggested that X-A10 domain substitutions, using a recombination site upstream to the C-A domain linker, performed better than C4-A10 domain substitutions. Overall, we consider that the ability to generate large library sizes more than compensates for the requirement to target conserved motif sequences. Even C1-A10 domain substitutions, which are most similar to early studies that kept C-A domains together and intact, were successful for metagenomic domain substitution. In the future we envisage that it might be possible to target more favourable recombination sites outside conserved motifs by amplifying and using large pools of known domains, and such hybrid approaches might prove to be very valuable in the long-term.

## Conclusions

We believe that NRPS domain substitution should embrace the stochastic nature of NRPS engineering, seeking to drown out the noise by turning up the volume. The field has aimed to develop clear and consistent rules for domain substitution.^5,12-14^ The difficulty to develop a ‘one size fits all’ approach emphasizes the advantages that a large-scale substitution approach can offer over lower throughput rational strategies. The success of C1-A10 domain substitution here is a pertinent example. Even though it was *ca*. 3-fold less effective than C4-A10 domain substitution, and the larger insert size is likely to be disadvantageous for a library generation perspective, it still provided access to many new peptide analogues. This suggests that our earlier work on C-A domain substitution^38^ would certainly have led to many new compounds, if only we had methods to create and screen large libraries as described in this study. Metagenomic domain substitution is likely applicable to many other non-ribosomal peptides as other papers have shown at least low levels of successful for domain substitution using at least one recombination site similar to our C4-A10 domain substitions.^15,17,32-35^ As has been shown by the field of directed evolution, sometimes the most rational approach is to accept the inherent noisiness and turn to semi-rational approaches that focus on volume.^4^

## Methods

### Calculating entropy and graphing average information over 18 bases

A combined codon-alignment containing NRPS modules for previously compiled sequences for *Pseudomonas* and *Streptomyces* species^15^ was generated in MUSCLE.^48^ This was processed using GBLOCK version 0.91b^49^ to remove regions of ambiguous alignment. Parameters for GBLOCK were: minimum number of sequences for a flank position set to 50% of total sequences, minimum block length of five, and gap positions permitted in up to half of the sequences. The python script “EntropyScanGraph.py” was used to calculate entropy and information according to Schneider and Stephens (1990). This was calculated for *Pseudomonas* and *Streptomyces* sequences both combined and separately. A graph was then generated showing the information averaged over 18 bp along the length of the alignment.

### Structural analysis

To identify a crystal structure with the highest percent identity and coverage for SCHEMA,^29^ the amino acid sequence of module 2 of PvdD was trimmed from the C1 to A10 motifs inclusive and submitted to the Swiss-Model server (http://swissmodel.expasy.org/). Structure 6P1J^30^ with 45.88% identity was selected as the top template. Sequence alignments PvdD module 2, 6P1J and eight alternate C-A sequences used as a source of A-domains (Supplementary Fig. 1) were generated using MUSCLE.^48^ SCHEMA was used to create a contact map and a python script used to calculate number of clashes introduced at each fusion point between Pa11 and the alternate modules. The PvdD module 2 trimmed sequence was submitted to the ThreaDomEx server (https://zhanggroup.org/ThreaDomEx/) to predict continuous and discontinuous domains.^31^ The structure 6P1J was examined and annotated in PyMOL (https://pymol.org/2/).

### DNA manipulation

All plasmids and sequences used in this study are provided in the Source Data file. Oligonucleotides were synthesised by Macrogen (Seoul, South Korea) and PCR amplification performed using Phusion polymerase (New England Biolabs; Ipswich, MA, USA) unless otherwise stated. Restriction enzymes were sourced from New England Biolabs. Solid and liquid media growth assays contained ampicillin (100 mg/L) and gentamycin (20 mg/L) for selection, unless otherwise specified.

For testing of conserved regions as recombination sites, constructs were created using the Gibson assembly-based NEBuilder Hifi DNA Assembly Master Mix (New England Biolabs) and transformed into *E. coli* DH5α. Purified plasmids were confirmed via Sanger sequencing (Macrogen) prior to transformation into a *P. aeruginosa ΔpvdD* strain.

For C1-A10 substitutions, the pDegC1V vector was created by modifying pUCBAD-SMC to contain a version of *pvdD* with the C1 to A10 motif region replaced by a linker containing silent restriction sites for SpeI and NotI. For C4-A10 substitutions, an equivalent vector pDegC4V was created in which the C4-A10 region of the second module of *pvdD* was replaced by a linker containing SpeI and SalI sites.

### Analysis of pyoverdine production

Strains were grown in 200 μL of low salt LB in a 96 well plate for 24 h at 37 °C, 200rpm, then 10 μL used to inoculate 190 μL of M9 media supplemented with 0.1 % (w/v) L-arabinose and 4 g/L succinate (pH 7.0). Following an additional 24 h incubation, cultures were pelleted and for each test well 100 μL of supernatant added to 100 μL of fresh M9 media in a new 96 well plate and absorbance measured at 400 nm using an EnSpire 2300 Multilabel Reader (PerkinElmer, Waltham, MA, USA). For mass spectrometry analysis, 1.5 μL of supernatant was mixed with 13.5 μL of matrix (500 μL acetonitrile, 500 μL ultrapure water, 1 μL trifluoroacetic acid, 10 μg α-cyano-4-hydroxycinnamic acid). Aliquots of 0.5 μL were spotted in triplicate onto an Opti-TOF® 384 well MALDI plate (Applied Biosystems, Foster City, CA) and allowed to dry at room temperature. Spots were analysed using a MALDI TOF/TOF 5800 mass spectrometer (Applied Biosystems) in positive ion mode. Peaks were externally calibrated using cal2 calibration mixture (Applied Biosystems) and labelled in Data Explorer (Applied Biosystems).

### Soil metagenomic DNA extraction and cosmid library construction

Metagenomic DNA was extracted from soil samples collected from Roxburgh (Central Otago, New Zealand) and St. Anne’s Lagoon (Cheviot, New Zealand) for the RX and SL metagenomic libraries, respectively. DNA was size selected, and cloned into the pWEB::tnc vector to create cosmid libraries in *E. coli* EC100 Δ*entD* cells as previously described (2022).^50^ Libraries were arrayed across multiple 96-well plates with each plate created separately. Titre samples from selected individual library wells were plated on LB agar plates containing ampicillin and chloramphenicol, with colony counts used for library size estimations. Cosmids were recovered from overnight cultures by miniprep.

### Degenerate primer design and testing

Sets of degenerate primers were designed to bind within motifs using HYDEN (Linhart & Shamir, 2005).^36^ Primers were 18 to 20 bp in length and differed in the level of degeneracy from 4 combinations to 128 combinations allowing 0 to 2 mismatches with the pooled *Pseudomonas* and *Streptomyces* sequences (Fig. 2). The resulting degenerate primer pairs were analysed with the Python script “PCR_screen.py” to count the number of sequences that contained binding sites while allowing for up to one mismatch.

The resulting 24 sets of C1-A10 primers and 12 sets of C4-A10 primers were tested by PCR amplification using Biomix Red™ reaction mastermix (Meridian Bioscience, Cincinnati, OH, USA). C1-A10 degenerate primer pairs were tested with eight random samples of SL metagenomic DNA using a standard PCR protocol which was then optimised for primer set 22 (95 °C 3 min; 35 cycles of: 95 °C 30s, 69 °C 30s, 72 °C 3 min; 72 °C 5 min). The twelve C4 forward primers were tested in conjunction with the 22 reverse primers with three samples of metagenomic DNA as template using a protocol that was then optimised for primer 10 with the 22-reverse primer (95 °C 2 min; 35 cycles of: 95 °C 30s, 63 °C 45s, 72 °C 2:45 min; 72 °C 10 min).

### Domain substitution library creation

To produce pools of NRPS domains for library creation, C1-A10 primers were adapted to contain an upstream SpeI and downstream NotI restriction site, and C4-A10 primers were modified to include a 25 bp homology region to pDegC4V to enable subsequent Gibson assembly. PCR was performed as above using C1-A10 primer set 22 or C4-A10 primer set 10, with metagenomic DNA as template and digested and gel-extracted C1-A10 pools or C4-A10 pools were ligated or assembled into pDegC1V or pDegC4V, respectively. The ligations and assemblies for each sub-library were used to transform *E. coli* DH5α. Following growth on solid media, colonies were scraped in 30% glycerol and combined in a centrifuge tube, followed by DNA extraction and purification. Titre samples were plated concurrently and used for colony counts to estimate library size.

### Sequence processing

To analyse library diversity and hits, the ends of the domain substitution inserts were Sanger sequenced in the forward and reverse direction using primers. End sequences were processed with the python script “Process_Sequencing.py” and those missing a degenerate primer site (allowing for 1 mismatch) were discarded as likely containing errors, as were too-short sequences containing vector on both ends of the insert, or sequences containing stop codons. The sequences were then trimmed starting from the 3’ end of the primer binding site to the maximum length that contained fewer than 0.5 expected errors based on Q scores from Sanger sequencing. Sequences containing stop codons or short sequences of less than 100 bp were discarded from further analysis. The OTUs were generated by clustering sequences at 95% identity using USEARCH 11.0.667. The longest read from each OTU was submitted to blastx (https://blast.ncbi.nlm.nih.gov/Blast.cgi) using the default parameters. For each OTU, the lowest E-value hit that had classification to the at least the phylum level was recorded. The R package iNEXT was used to estimate OTU diversity.^37^

### Library screening

Pilot screening of the C1-A10 libraries was carried out using a dilution strategy. C1-A10 sub-libraries were used to transform *P. aeruginosa* PAO1 Δ*pvdD*, which was then diluted to a concentration of ∼5 cells/well in 200 μL LB broth. Pyoverdine production was assessed for duplicates of each well using a MALDI TOF/TOF 5800 mass spectrometer (Applied Biosystems). Wells from which a unique pyoverdine analogue were detected were streaked on solid media followed by testing to isolate the pyoverdine-producing strains.

For colony-based screening, C1-A10 and C4-A10 sub-libraries were used to transform *P. aeruginosa* PAO1 Δ*pvdD* which was plated on solid media. Colonies were then inoculated into three 96 well plates per sub-library (4608 colonies in total). After 24 h growth at 37 °C, 200rpm, 10 μL of culture was used to inoculate 190 μL of M9 media supplemented with 0.1 % (w/v) L-arabinose and 4 g/L succinate (pH 7.0). Cultures were grown for 37 °C for 24 h, 200rpm after which, emission at 440 nm was measured using an EnSpire 2300 Multilabel Reader (PerkinElmer). Wells which had a fluorescence reading at 8000 or above (approx. 4% of WT fluorescence) were selected for retesting using the analysis of pyoverdine production protocol described above. Data was processed with python scripts and R was used to create graphs.

### Extraction and LCMS analysis of pyoverdine

After 16 h growth of each test strain in low salt LB at 37 °C, 200rpm, 1 mL of culture was used to inoculate 24 mL of M9 media supplemented with 0.1 % (w/v) L-arabinose, 1% casamino acids and 4 g/L succinate (pH 7.0) in 50 mL shaking flasks. After 24 h growth at 37 °C, 200 rpm, cultures were centrifuged and supernatant loaded onto activated HLB columns (CHROMABOND, MACHEREY-NAGEL, Düren, Germany) on vacuum. Columns were washed with 0.5 mL double distilled water and eluted in 2 mL 100% Methanol. Solvent was removed under vacuum and the residue resuspended in 100 μL 50% acetonitrile. For HPLC-MS analysis an Agilent 6530 Accurate-Mass Q-TOF LC/MS system equipped with an Agilent 1260 HPLC system (Agilent, Santa Clara, CA) was used. The mobile phase consisted of water + 0.1% formic acid (solvent A) and acetonitrile + 0.1% formic acid (solvent B). Extracts were passed through an Eclipse Plus C18, 3.5 μm, 2.1 x 30mm, column (Agilent) heated to 40 °C at a flow rate of 0.4 ml/min over 8 minutes using a gradient of 3%-60% solvent B. The spray voltage was 3.50 kV, gas temperature was 300 °C, and collision energy was 50 eV.

Mass spectrometry data files for each pyoverdine sample were visualised in MassHunter Qualitative Analysis Navigator B.08.00 followed by processing using MZmine 3.^51^ MS and MS/MS data was extracted and chromatograms were built. ^13^C isotopes were filtered out and duplicate peaks within a 2 minute retention time window were merged. A molecular network was created using the online workflow (https://ccms-ucsd.github.io/GNPSDocumentation/) on the GNPS website (http://gnps.ucsd.edu).^44^ Edges were filtered to have a cosine score above 0.5 and more than 3 matched peaks and were kept in the network only if each of the nodes appeared in each other’s respective top 12 most similar nodes. The molecular network was visualised in Cytoscape version 3.8.0.^52,53^ Node size was set to peak area of extracted ion chromatograms for each detected m/z in the network, and normalised across the different pyoverdine extractions by setting the highest abundance peak for each to 1000000. Edge width was set to the cosine value. Nodes were annotated based on known pyoverdine adducts and side chain variations,^39,40^ including chelation of aluminium.^41^

## Data availability

Source data are provided with this paper.

## Code availability

Code generated during this study is available on GitHub via the following link: https://github.com/MarkCalcott/metagenomic-domain

## Acknowledgements

This work was supported by the Royal Society of New Zealand Marsden Fund (Grant 18-VUW-082 to M.J.C. and G.L.C), and the Health Research Council of New Zealand (contract 16/172 to D.F.A. and J.G.O.). S.R.M. was supported by a New Zealand Lottery Health Research Scholarship (contract LHR-2020-129624).

## Contributions

S.R.M., D.F.A. and M.J.C. conceived and designed the work. S.R.M., E.M.M and M.J.C. performed the experiments. S.R.M. and M.J.C. performed the data analysis and prepared all figures. L.J.S, J.G.O. and M.J.C created the SL and RX metagenomic libraries. M.J.C., G.L.C, D.F.A and J.G.O. obtained funding for this study. S.R.M., D.F.A. and M.J.C. wrote the manuscript, and all authors approved the final version.

