## Supplementary materials for "Metagenomic domain substitution for the high-throughput modification of non-ribosomal peptide analogues"

**Supplementary Information:**

Supplementary Figures 1 – 10

Supplementary Tables 1 – 5

**Supplementary Figure 1. Amino acid sequence alignment of PvdD M2 and the eight alternative C-A domain sequences used for SCHEMA analysis and substitution testing.** Sequences are aligned from the C1 to A10 motifs inclusive. Conserved sequence motifs and recombination sites, including X, 1 and 2, are labelled. Image created using Geneious version 8.1 (Biomatters. Available from http://www.geneious.com). Continued on the following page.

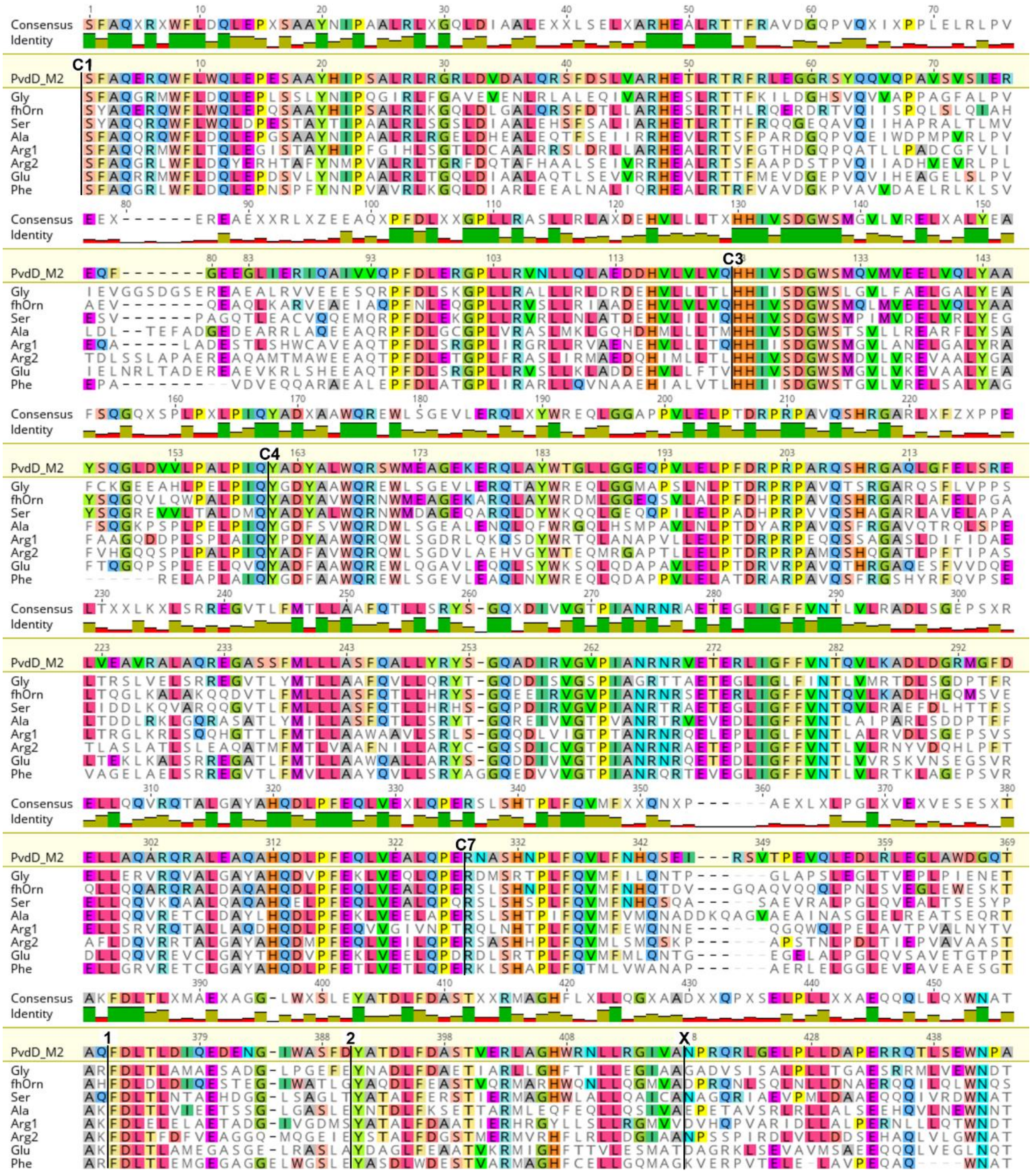

### Supplementary Figure 1 continued

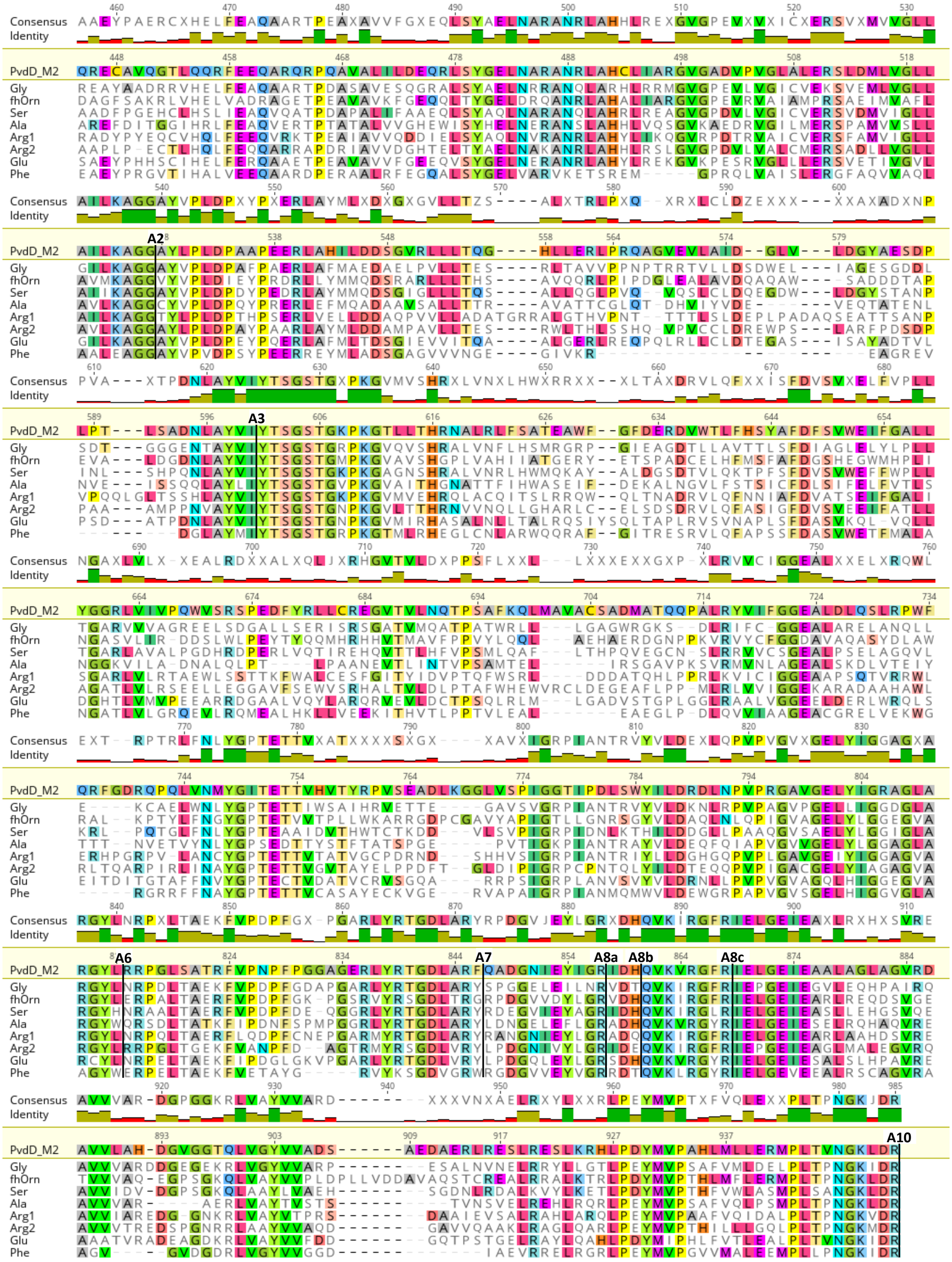

**Supplementary Figure 2. Mass spectra for pyoverdine variants from recombination site testing.** MALDI-TOF mass spectra from *P. aeruginosa* PAO1 *pvdD* deletion strains expressing *pvdD* constructs bearing substitutions of alternate domains using **a.** the C1-A10 recombination boundaries, or **b.** C4-A10 recombination boundaries. Labels indicate the *m/z* ratio for $[M+H]^+$ ions corresponding to pyoverdine with a succinamide acyl moiety in which the terminal L-Thr residue has been replaced with another amino acid. A total of n = 3 independent experiments were performed with consistent results; representative spectra are presented here. Continued on the following page.

**a.** Mass spectra from C1-A10 substitutions

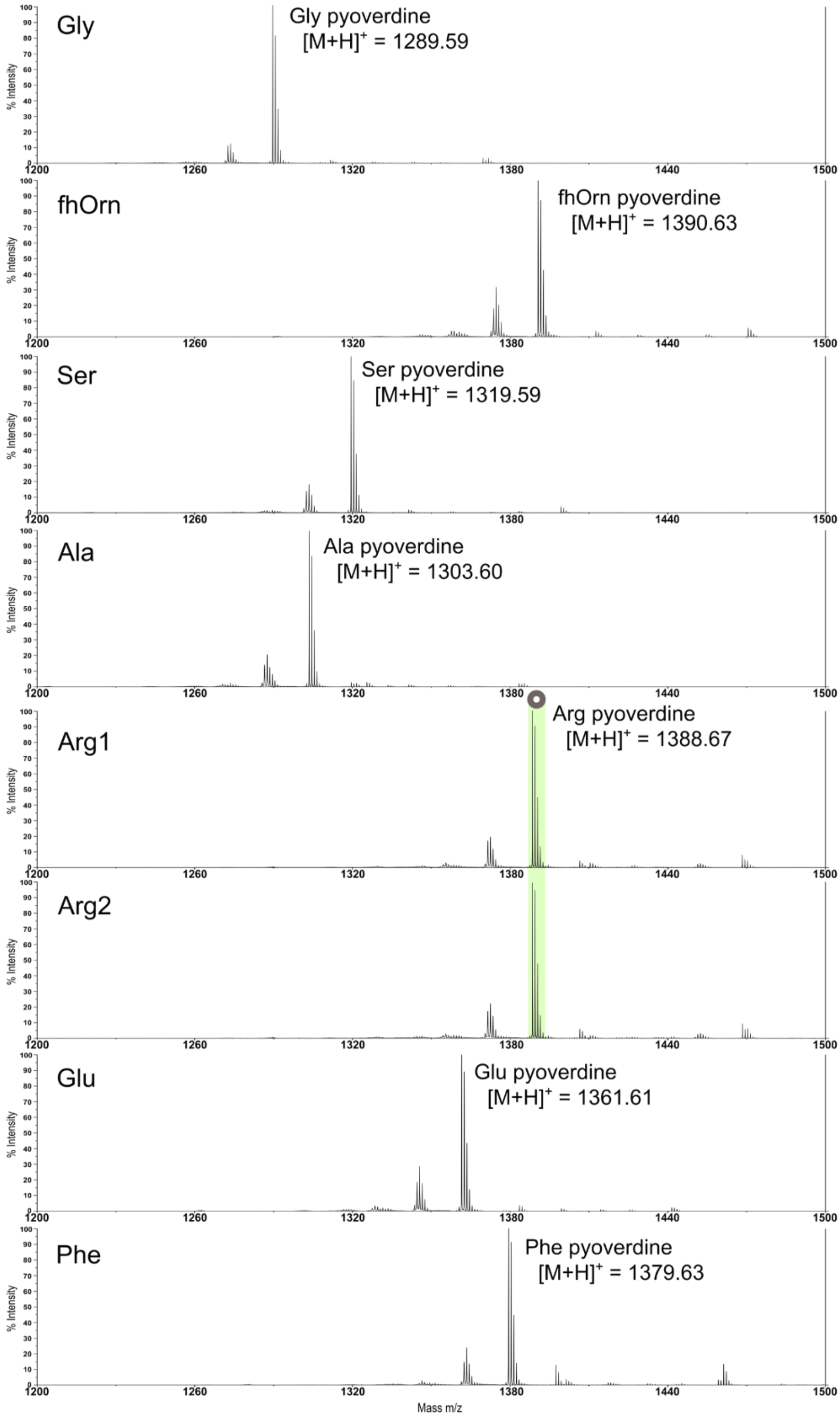

**Supplementary Figure 2 continued**

**b.** Mass spectra from C4-A10 substitutions

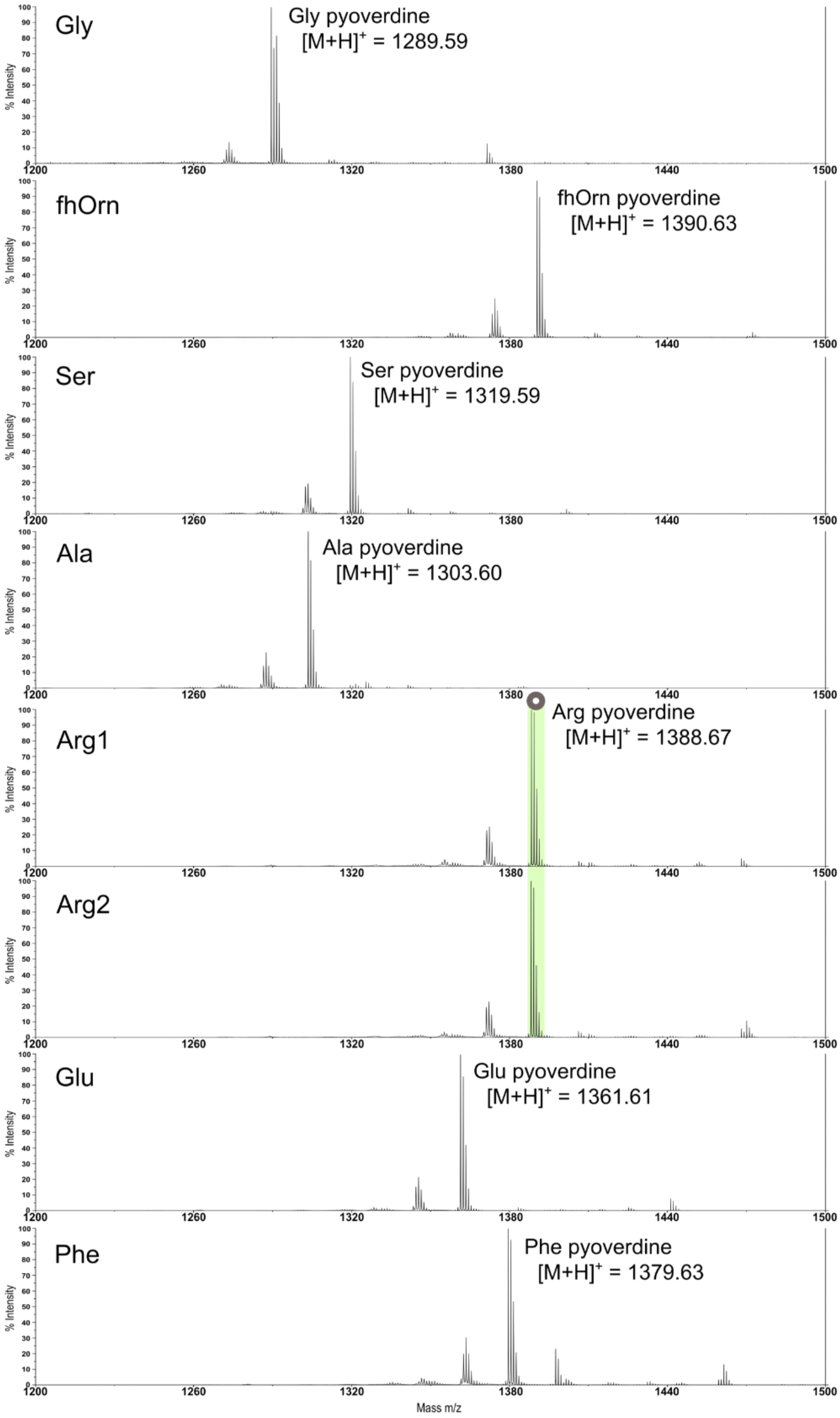

**Supplementary Figure 3. Testing of 24 sets of C1-A10 degenerate primers.** Primers were tested using a set PCR protocol with eight samples of SL metagenomic DNA as template. The gel photo shows amplification for all 24 primer sets, labelled 1 to 24, and the expected product band size of 3,000 bp is indicated.

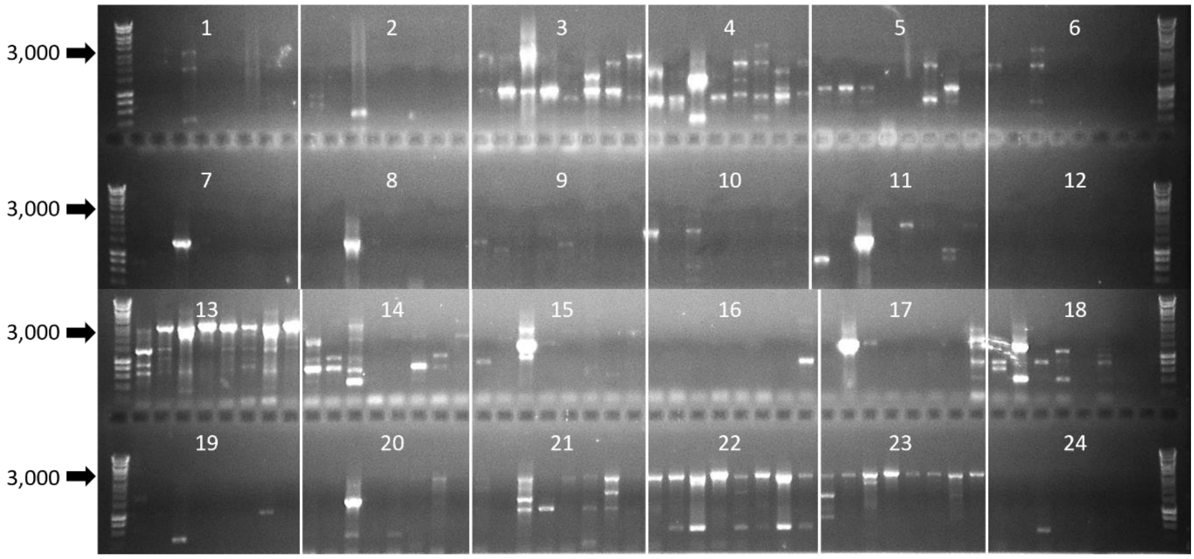


**Supplementary Figure 4. Testing of 12 sets of C4 forward degenerate primers. a.** Twelve sets of C4-A10 degenerate primers (labelled 1-12) were tested as per Supplementary Figure 3 except with three samples of SL or RX metagenomic DNA as template. **b.** Expanded testing of primer set 10 using eight samples of SL metagenomic DNA as template. The expected product band size of 2,400 bp is indicated.

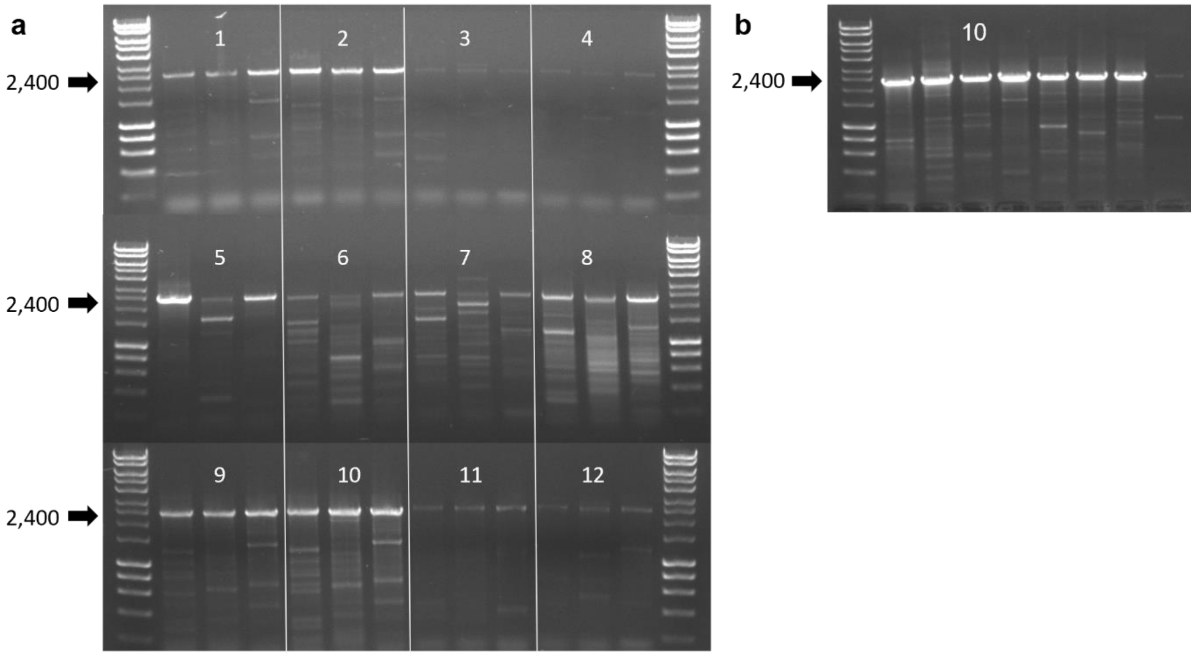

**Supplementary Figure 5. C1-A10 PCR amplification from individual wells of eight RX1-RX4 and SL1-SL4 metagenomic sub-libraries arrayed in 96 well plates.** PCR was performed in 96 well plates using C1-A10 primer set 22 and metagenomic DNA sub-libraries RX1-RX1 and SL1-SL4. The expected product band size of 3,000 bp is indicated.

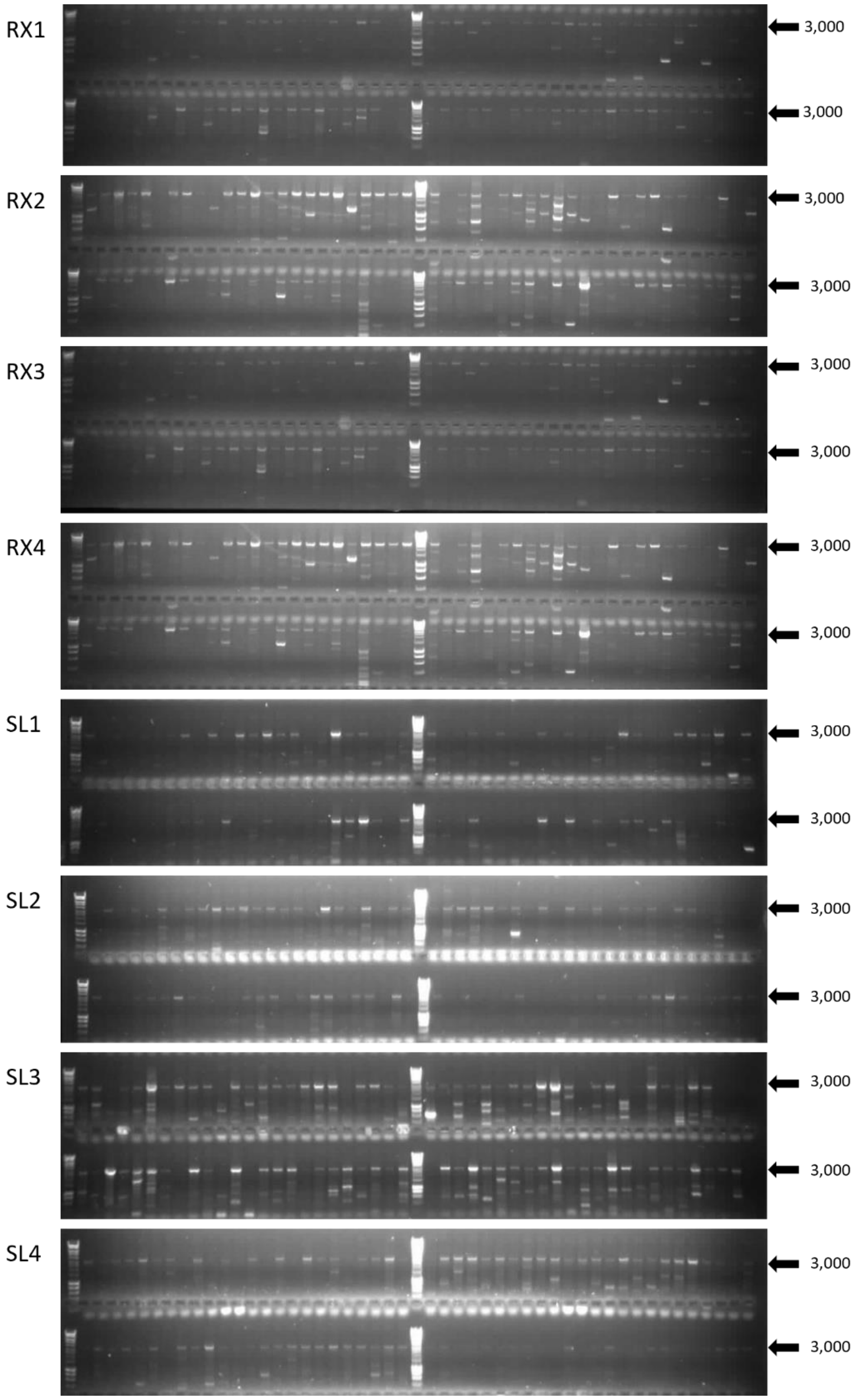

**Supplementary Figure 6. C4-A10 PCR amplification from individual wells of eight metagenomic DNA template sets from the arrayed sub-libraries RX1-RX4 and SL1-SL4.** Amplification as per Supplementary Figure 5 except using C4-A10 primer set 10.

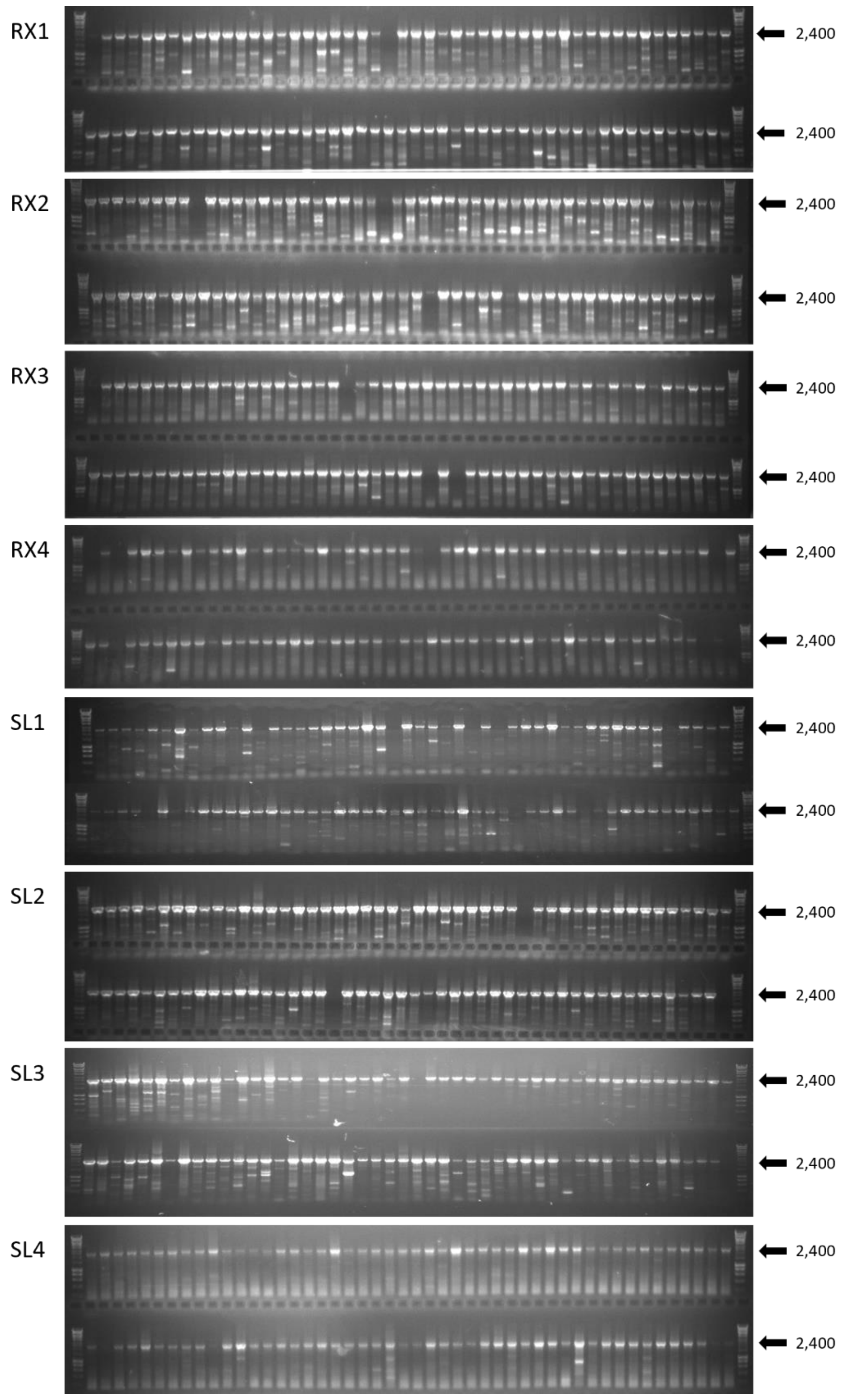

**Supplementary Figure 7. Activity for the pyoverdine variants detected from preliminary screening of the C1-A10 libraries.** Standard amino acid abbreviations are used, with numbering distinguishing alternative modules with the same specificity. In all cases, n = 6 independent experiments and data are presented as mean values ± SD. Source data are provided as a Source Data file.

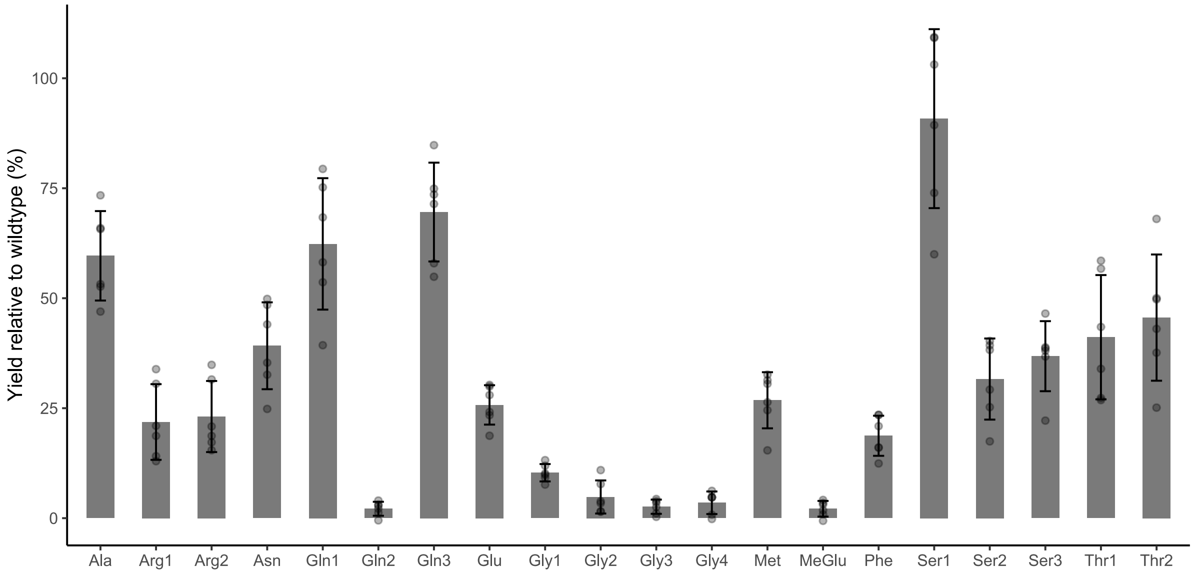

**Supplementary Figure 8. MALDI-TOF mass spectra for the pyoverdine variants detected from preliminary screening of the C1-A10 libraries.** The *m/z* ratio of $[M+H]^+$ ions corresponding to pyoverdine with a succinamide acyl moiety are labelled. Standard abbreviation codes are used to identify the amino acids, with methyl-Glu to indicate a m/z ratio consistent with methylated Glu. A total of n= 3 independent experiments were performed with consistent results; representative spectra are presented here.

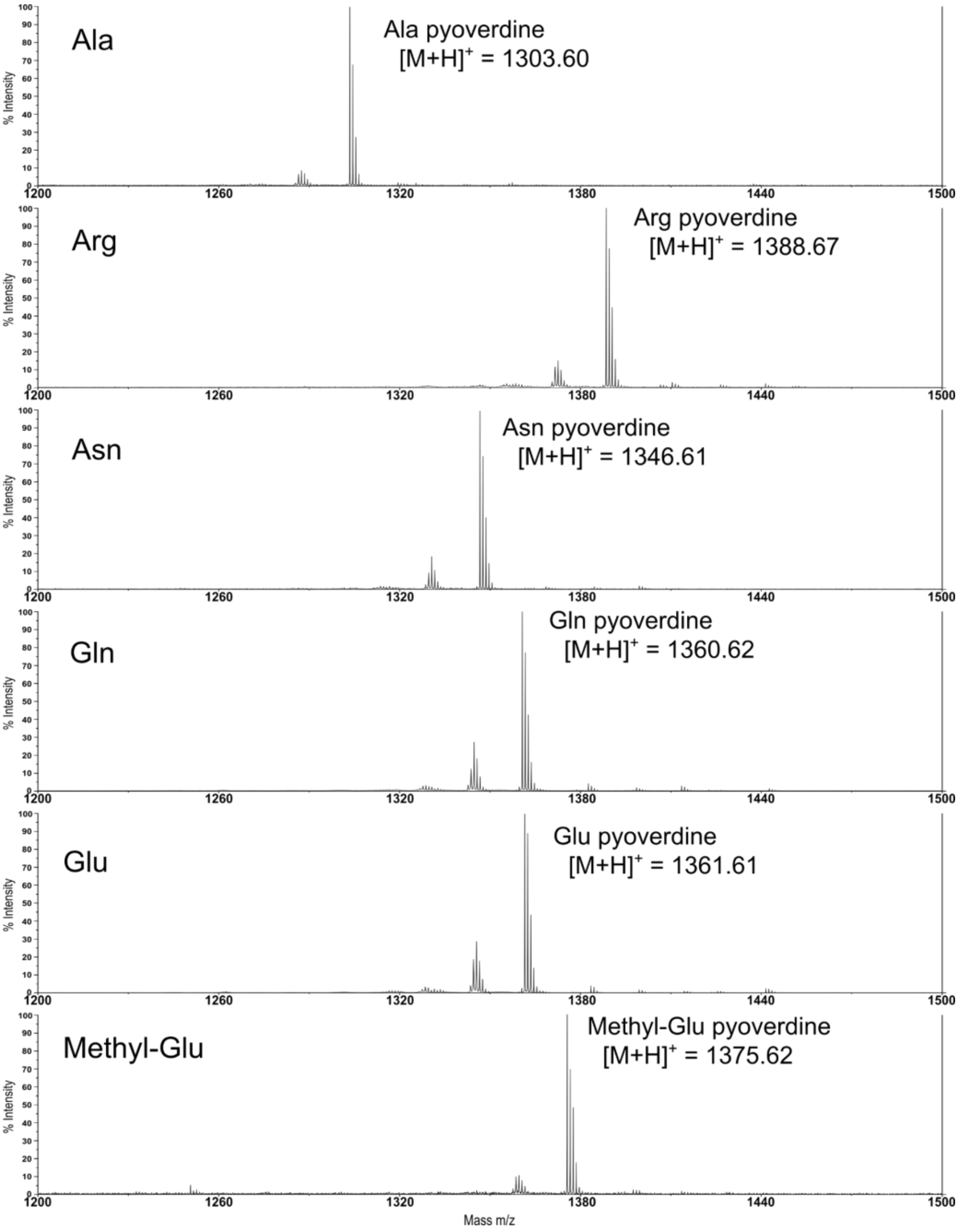


**Supplementary Figure 8 continued**

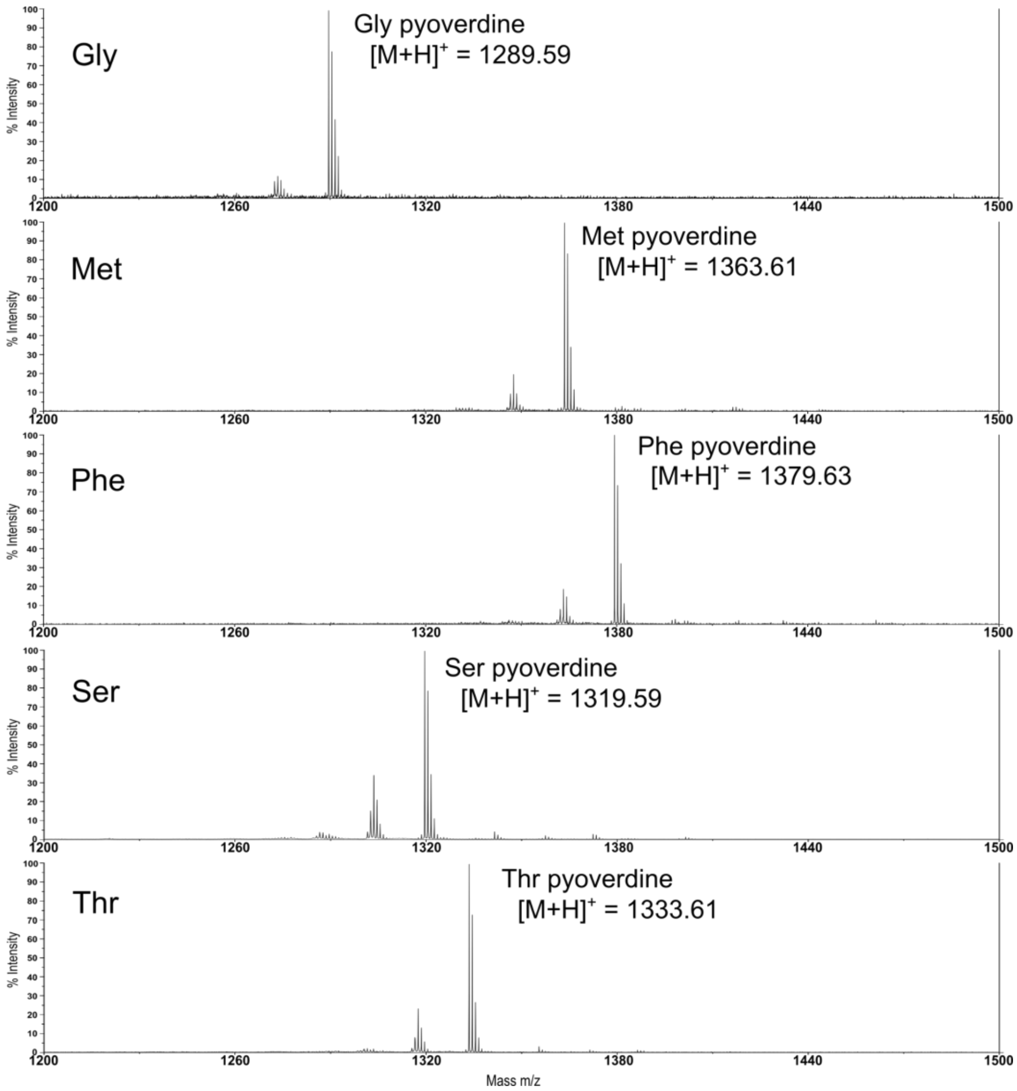
Gly
Gly pyoverdine
[M+H]+ = 1289.59
Met
Met pyoverdine
[M+H]+ = 1363.61
Phe
Phe pyoverdine
[M+H]+ = 1379.63
Ser
Ser pyoverdine
[M+H]+ = 1319.59
Thr
Thr pyoverdine
[M+H]+ = 1333.61
% Intensity
Mass m/z
1200
1260
1320
1380
1440
1500

**Supplementary Figure 9. MALDI-TOF mass spectra for the pyoverdine variants detected from colony-based screening of the C1-A10 and C4-A10 libraries.** Spectra were obtained from the top yielding strains producing each substituted pyoverdine variant. The *m/z* ratio of $[M+H]^+$ ions corresponding to pyoverdine with a succinamide acyl moiety are labelled. Standard abbreviation codes are used to identify the amino acids, with fhOrn designating N5-formyl-N5-hydroxyornithine and Xle designating Leu or Ile. A total of n = 3 independent experiments were performed with consistent results; representative spectra are presented here.

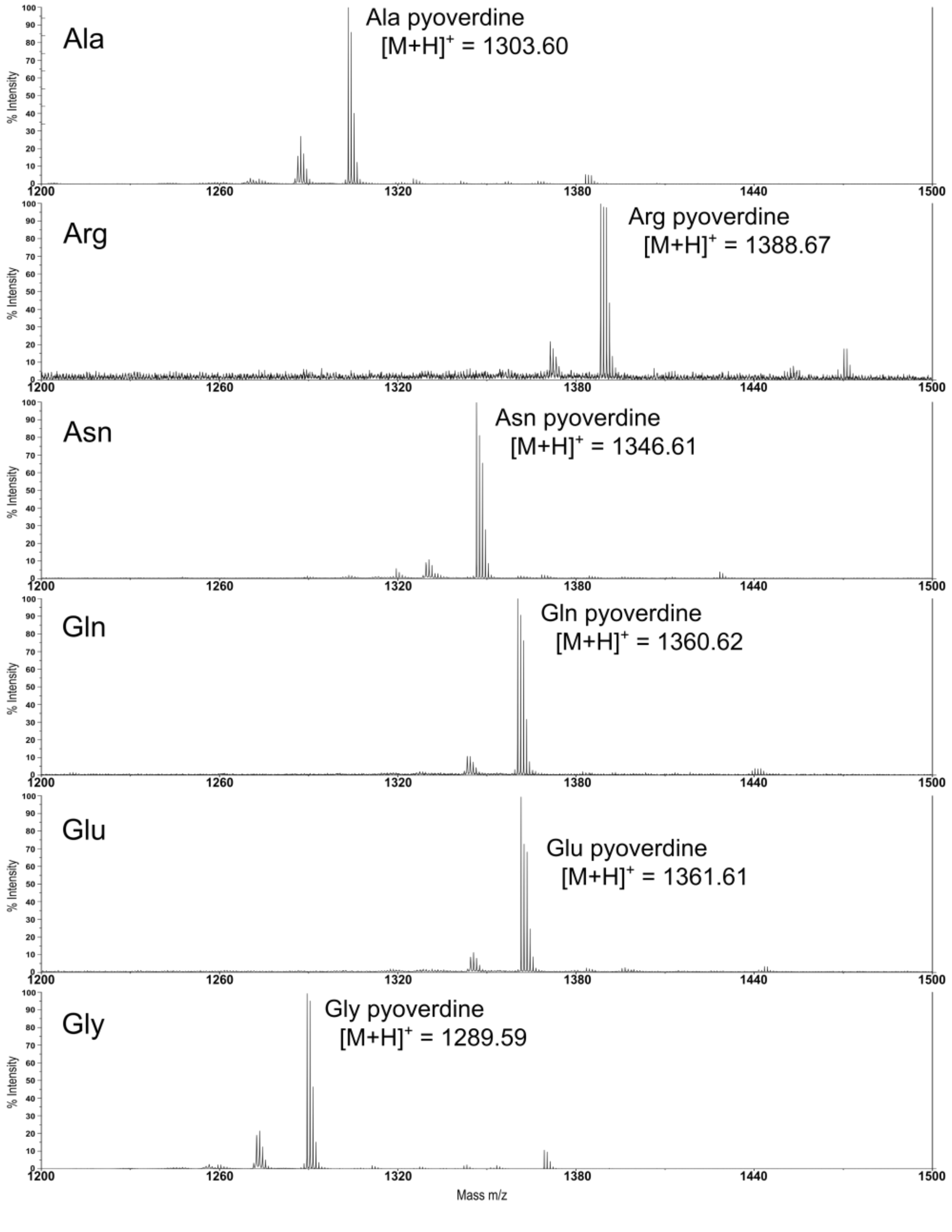

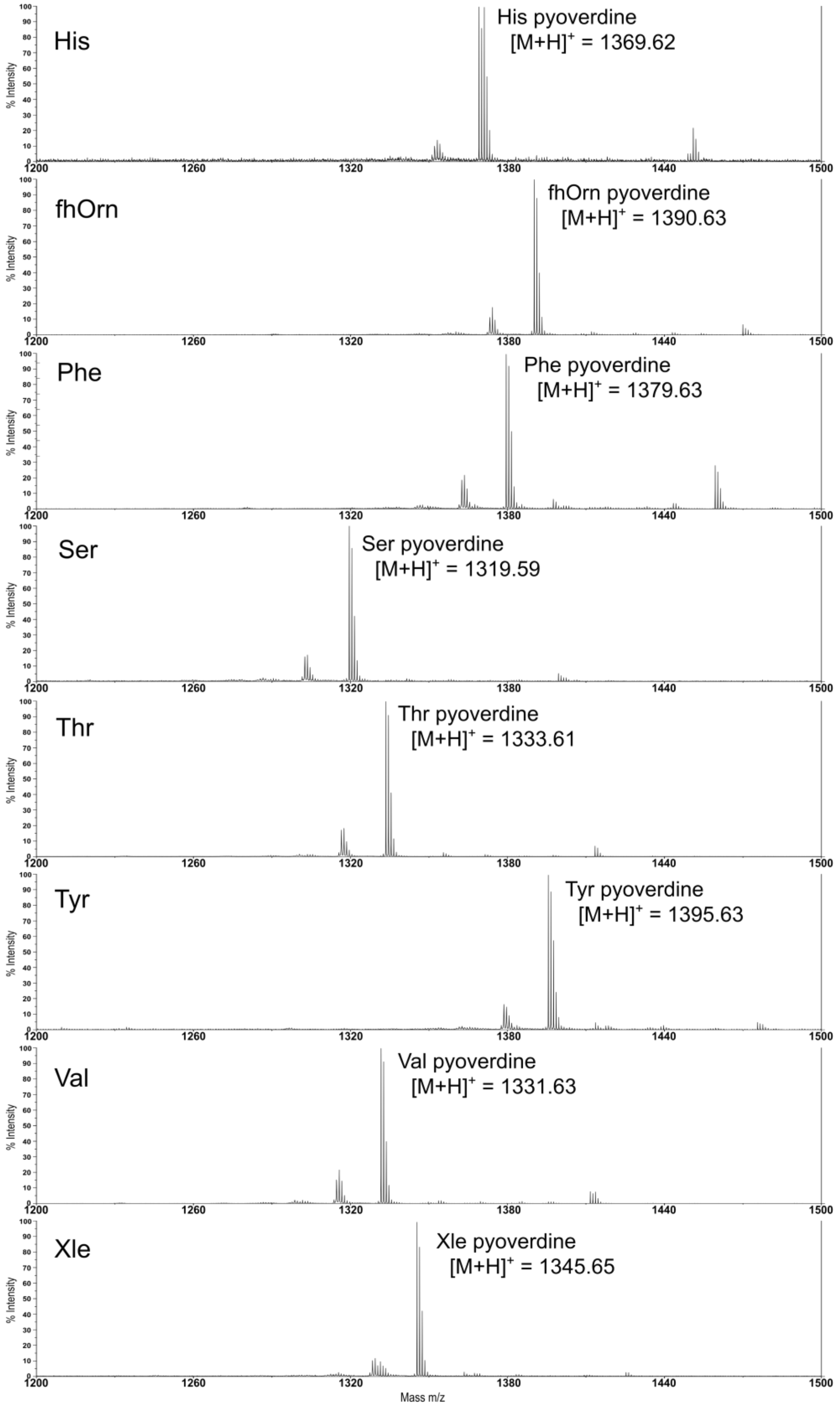

His
His pyoverdine
[M+H]+ = 1369.62
fhOrn
fhOrn pyoverdine
[M+H]+ = 1390.63
Phe
Phe pyoverdine
[M+H]+ = 1379.63
Ser
Ser pyoverdine
[M+H]+ = 1319.59
Thr
Thr pyoverdine
[M+H]+ = 1333.61
Tyr
Tyr pyoverdine
[M+H]+ = 1395.63
Val
Val pyoverdine
[M+H]+ = 1331.63
Xle
Xle pyoverdine
[M+H]+ = 1345.65
% Intensity
1200 1260 1320 1380 1440 1500
Mass m/z

**Supplementary Figure 10. Molecular networks of the highest yielding strains for each of the pyoverdine variants detected from colony-based screening of the C1-A10 and C4-A10 libraries.** LC-MS data was obtained from the top yielding strains producing each substituted pyoverdine variant and analysed using Mzmine 3, GNPS and Cytoscape version 3.8.0 . Node size is area under the curve of extracted ion chromatograms for each detected m/z in the network, and normalised across the different pyoverdine extractions by setting the highest abundance peak for each to 1000000. Edge width is the cosine value to reflect the relatedness of the nodes. A table of node data, peak area and annotations for the networks is in Supplementary Table 4.

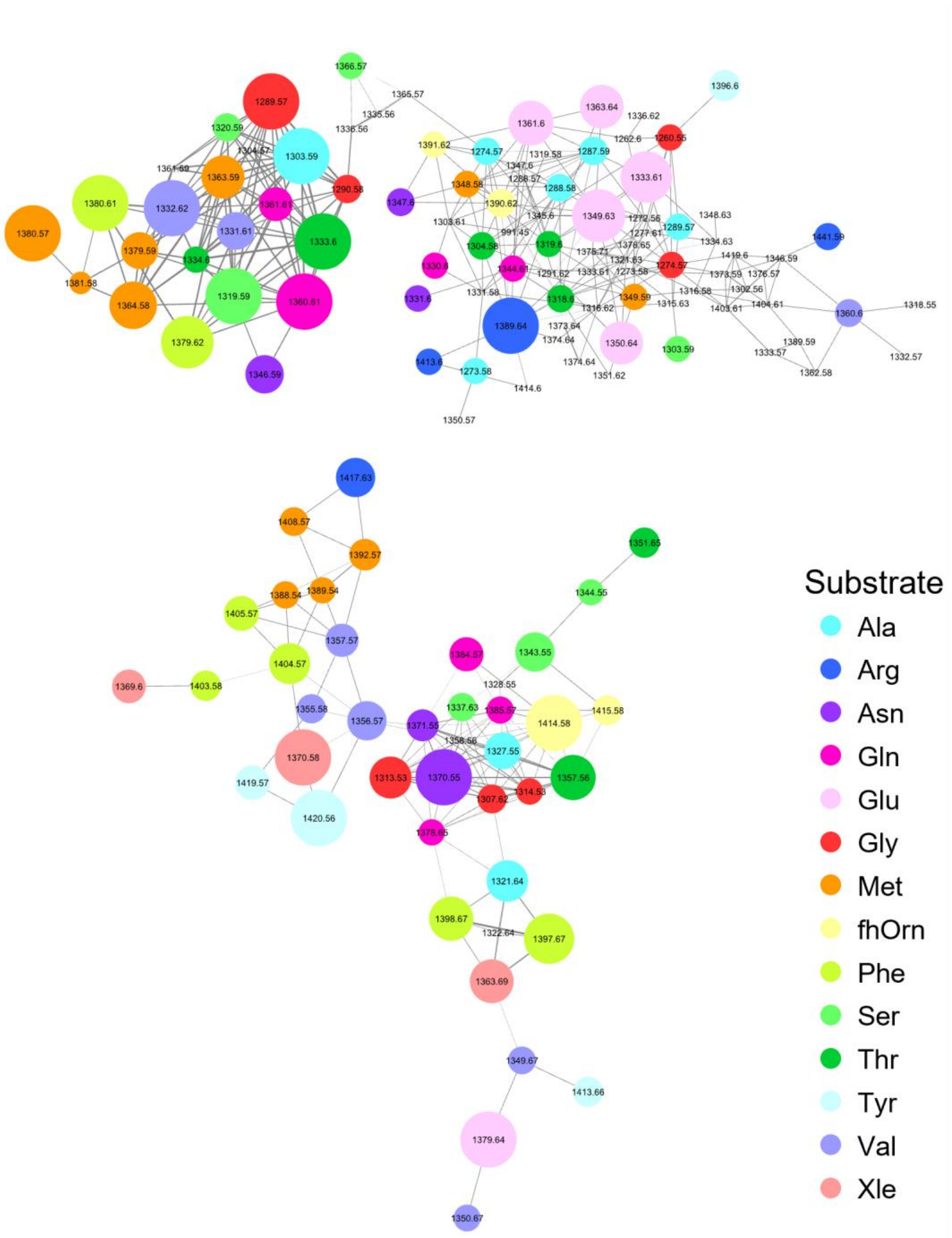

**Supplementary Table 1. Degenerate primers tested in this study and the percentage of *Pseudomonas*, *Streptomyces* and *Bacillus* DNA sequences that contained binding sites.** DNA sequences were of C-A-T domains and collated as part of a previous study.[15] Binding sites were identified allowing for 1 mismatch per primer. Primers are labelled as either C1-A10 or C4-A10 depending on the region they amplify followed by the primer set number. Primer sets used for library building are indicated in bold.

| Primer set | Forward primer degeneracy | Reverse primer degeneracy | *Pseudomonas* | *Streptomyces* | *Bacillus* |
|---|---|---|---|---|---|
| C1-A10_1 | 4 | 4 | 14.6% | 5.6% | 0.0% |
| C1-A10_2 | 4 | 4 | 10.8% | 0.0% | 0.0% |
| C1-A10_3 | 4 | 4 | 14.4% | 2.8% | 0.0% |
| C1-A10_4 | 16 | 16 | 14.0% | 5.2% | 0.0% |
| C1-A10_5 | 16 | 16 | 13.5% | 14.6% | 0.0% |
| C1-A10_6 | 16 | 16 | 16.2% | 8.5% | 0.0% |
| C1-A10_7 | 8 | 16 | 19.9% | 1.4% | 0.0% |
| C1-A10_8 | 16 | 16 | 23.8% | 8.5% | 0.0% |
| C1-A10_9 | 32 | 24 | 24.3% | 3.3% | 0.0% |
| C1-A10_10 | 32 | 32 | 29.3% | 9.4% | 0.0% |
| C1-A10_11 | 32 | 32 | 25.6% | 8.0% | 0.0% |
| C1-A10_12 | 32 | 32 | 31.8% | 13.1% | 0.0% |
| C1-A10_13 | 64 | 32 | 4.3% | 24.9% | 0.0% |
| C1-A10_14 | 64 | 64 | 26.8% | 18.8% | 0.0% |
| C1-A10_15 | 32 | 64 | 35.5% | 5.6% | 0.0% |
| C1-A10_16 | 64 | 64 | 37.3% | 9.9% | 0.0% |
| C1-A10_17 | 64 | 64 | 34.3% | 8.0% | 0.0% |
| C1-A10_18 | 64 | 64 | 32.3% | 10.3% | 0.0% |
| C1-A10_19 | 64 | 64 | 34.3% | 18.8% | 0.0% |
| C1-A10_20 | 64 | 32 | 33.0% | 17.4% | 0.0% |
| C1-A10_21 | 128 | 54 | 36.2% | 24.9% | 0.0% |
| **C1-A10_22** | **96** | **54** | **38.4%** | **42.7%** | **0.0%** |
| C1-A10_23 | 128 | 64 | 14.9% | 29.6% | 0.0% |
| C1-A10_24 | 128 | 128 | 40.3% | 16.9% | 0.0% |
| C4-A10_1 | 128 | 54 | 41.0% | 44.1% | 1.4% |
| C4-A10_2 | 96 | 54 | 38.2% | 38.0% | 0.8% |
| C4-A10_3 | 64 | 54 | 41.4% | 37.6% | 0.3% |
| C4-A10_4 | 64 | 54 | 37.5% | 41.3% | 0.0% |
| C4-A10_5 | 32 | 54 | 31.8% | 37.1% | 0.0% |
| C4-A10_6 | 128 | 54 | 26.8% | 24.4% | 0.0% |
| C4-A10_7 | 128 | 54 | 24.5% | 17.8% | 0.8% |
| C4-A10_8 | 128 | 54 | 40.7% | 36.6% | 0.0% |
| C4-A10_9 | 128 | 54 | 41.6% | 38.0% | 2.7% |
| **C4-A10_10** | **128** | **54** | **38.9%** | **45.1%** | **2.7%** |
| C4-A10_11 | 64 | 54 | 43.5% | 37.6% | 0.0% |
| C4-A10_12 | 64 | 54 | 40.0% | 35.7% | 0.0% |

**Supplementary Table 2. Library sizes for each sub-library of the RX and SL metagenomic libraries.** The RX and SL metagenomic libraries were each stored in four sub-libraries labelled RX1-RX4 and SL1-SL4, with each sub-library containing 96-wells. Unique clones were extrapolated from colony counts taken from a sample during library creation.

| Metagenomic library | Sub-library | Unique clones | Clones per well |
|---|---|---|---|
| Roxburgh (RX) | RX1 | 1,750,000 | 18,229 |
| | RX2 | 1,080,000 | 11,250 |
| | RX3 | 3,570,000 | 37,188 |
| | RX4 | 2,090,000 | 21,771 |
| St. Anne's Lagoon (SL) | SL1 | 1,130,000 | 11,771 |
| | SL2 | 4,440,000 | 46,250 |
| | SL3 | 2,900,000 | 30,208 |
| | SL4 | 4,430,000 | 46,146 |

**Supplementary Table 3. Library size for each metagenomic domain substitution sub-library.** Metagenomic library and sub-library indicates the source DNA from Supplementary Table 2 used for degenerate PCR reactions. Unique clones were extrapolated from colony counts taken from a sample during library creation.

| Metagenomic library | Substitution type | Sub-library | Unique clones |
|---|---|---|---|
| Roxburgh (RX) | C1-A10 | RX1 | 1,850 |
| | | RX2 | 4,700 |
| | | RX3 | 3,100 |
| | | RX4 | 2,200 |
| | C4-A10 | RX1 | 52,000 |
| | | RX2 | 50,000 |
| | | RX3 | 25,000 |
| | | RX4 | 150,000 |
| St. Anne's Lagoon (SL) | C1-A10 | SL1 | 4,600 |
| | | SL2 | 2,200 |
| | | SL3 | 7,500 |
| | | SL4 | 1,800 |
| | C4-A10 | SL1 | 57,000 |
| | | SL2 | 50,000 |
| | | SL3 | 35,000 |
| | | SL4 | 160,000 |

**Supplementary Table 4. Annotated unique pyoverdine nodes from the molecular networking of the highest yielding strains for each of the sixteen pyoverdine variants (Supplementary Figure 10).** Parent m/z indicates the detected parent mass as labelled on the molecular networks. Peak area was calculated from the peak area of extracted ion chromatograms of the detected parent m/z, which was then normalised for each substrate by setting the highest abundance ion to 1,000,000. Annotations were made based on the theoretical m/z of plausible pyoverdine derivatives.

| Parent m/z | Substrate | Peak area | Normalised Peak area | Annotation | Theoretical m/z | ppm |
|---|---|---|---|---|---|---|
| 1303.59 | Ala | 7475739 | 1000000 | Ala succinamide | 1303.603 | 9.63 |
| 1304.57 | Ala | 235970.9 | 31564.9 | Ala succinate | 1304.587 | 9.63 |
| 1332.57 | Ala | 430121.8 | 57535.7 | Ala α-Ketoglutaric acid | 1332.581 | 8.61 |
| 1441.59 | Arg | 609927.2 | 92163.96 | Arg succinamide + Fe | 1441.578 | -8.31 |
| 1389.64 | Arg | 6617850 | 1000000 | Arg succinate | 1389.651 | 7.60 |
| 1417.63 | Arg | 3124707 | 472163.5 | Arg α-Ketoglutaric acid | 1417.645 | 10.92 |
| 1346.59 | Asn | 1876980 | 443574.3 | Asn succinamide | 1346.608 | 13.64 |
| 1347.6 | Asn | 498636.1 | 117839.3 | Asn succinate | 1347.592 | -5.65 |
| 1390.62 | fhorn | 495563 | 138173.4 | fhOrn succinamide | 1390.635 | 10.49 |
| 1391.62 | fhorn | 228969.5 | 63841.54 | fhOrn succinate | 1391.619 | -1.01 |
| 1419.6 | fhorn | 189222.6 | 52759.26 | fhOrn α-Ketoglutaric acid | 1419.614 | 8.11 |
| 1360.61 | Gln | 8725238 | 1000000 | Gln succinamide | 1360.624 | 10.30 |
| 1361.61 | Gln | 2673065 | 306360.1 | Gln succinate | 1361.608 | -1.44 |
| 1389.59 | Gln | 396301.9 | 45420.18 | Gln α-Ketoglutaric acid | 1389.603 | 9.32 |
| 1361.6 | Glu | 109774.8 | 648491.7 | Glu succinamide | 1361.608 | 5.90 |
| 1289.57 | Gly | 6860081 | 1000000 | Gly succinamide | 1289.587 | 13.10 |
| 1290.58 | Gly | 923116.1 | 134563.5 | Gly succinate | 1290.571 | -7.04 |
| 1318.55 | Gly | 347202.7 | 50612.05 | Gly α-Ketoglutaric acid | 1318.566 | 12.00 |
| 1380.57 | Met | 3678745 | 1000000 | Met Malic acid | 1380.587 | 12.26 |
| 1379.59 | Met | 1458598 | 396493.4 | Met Malic amide | 1379.603 | 9.36 |
| 1363.59 | Met | 2161310 | 587513 | Met succinamide | 1363.608 | 13.20 |
| 1364.58 | Met | 2668132 | 725283.3 | Met succinate | 1364.592 | 8.81 |
| 1392.57 | Met | 856407.2 | 232798.8 | Met α-Ketoglutaric acid | 1392.587 | 12.16 |
| 1379.62 | Phe | 1325042 | 885610.8 | Phe succinamide | 1379.634 | 10.04 |
| 1380.61 | Phe | 1496191 | 1000000 | Phe succinate | 1380.618 | 5.70 |
| 1319.59 | Ser | 3101965 | 1000000 | Ser succinamide | 1319.597 | 5.57 |
| 1320.59 | Ser | 396706.9 | 127888.9 | Ser succinate | 1320.581 | -6.54 |
| 1333.6 | Thr | 6354238 | 1000000 | Thr succinamide | 1333.613 | 9.84 |
| 1334.6 | Thr | 443770.2 | 69838.46 | Thr succinate | 1334.597 | -2.15 |
| 1362.58 | Thr | 310813.6 | 48914.38 | Thr α-Ketoglutaric acid | 1362.592 | 8.84 |
| 1419.57 | Tyr | 193023.7 | 324078.2 | Tyr succinamide +Al | 1419.587 | 11.86 |
| 1396.6 | Tyr | 133279 | 223769.6 | Tyr succinate | 1396.613 | 9.15 |
| 1331.61 | Val | 886126.9 | 422473.8 | Val succinamide | 1331.634 | 17.91 |
| 1332.62 | Val | 2097472 | 1000000 | Val succinate | 1332.618 | -1.60 |
| 1360.6 | Val | 248072.2 | 118272 | Val α-Ketoglutaric acid | 1360.613 | 9.39 |
| 1369.6 | Xle | 152882.9 | 299485 | Xle succinamide +Al | 1369.608 | 5.52 |

| | | | | | | |
|---|---|---|---|---|---|---|
| 1370.58 | Xle | 510486 | 1000000 | Xle succinate +Al | 1370.592 | 8.45 |
| 1393.566 | His | 829888.8 | 1000000 | His succinamide + Al | 1393.582 | 11.78 |
| 1394.554 | His | 558511 | 672995 | His succinate +Al | 1394.566 | 8.91 |
| 1399.566 | MeGlu | 46241.63 | 1000000 | Me-Glu succinamide + Al | 1399.582 | 11.48 |
